# Evolution of NLR resistance genes with non-canonical N-terminal domains in wild tomato species

**DOI:** 10.1101/786194

**Authors:** Kyungyong Seong, Eunyoung Seo, Meng Li, Brian Staskawicz

**Author notes:** These authors contributed equally to this work (Kyungyong Seong, Eunyoung Seo).

## Abstract

**Background:** Nucleotide-binding and leucine-rich repeat immune receptors (NLRs) are an important component of plant immunity that provides resistance against diverse pathogens. NLRs often exist as large gene families, the members of which display diverse multi-domain architectures (MDAs) and evolve through various mechanisms of duplications and selections.

**Results:** We conducted resistance gene enrichment sequencing (RenSeq) with single-molecule real time (SMRT) sequencing of PacBio for 18 accessions in Solanaceae including 15 wild tomatoes. We demonstrate what was previously known as Solanaceae Domain (SD) not only is more diverse in structure and function but also far anciently originated from the most recent common ancestor (MRCA) between Asterids and Amaranthaceae. In tomato, NLRs with the extended N-terminus displayed distinct patterns of evolution based on phylogenetic clades by proliferation, continuous elongation and domain losses.

**Conclusions:** Our study provides high quality gene models of NLRs that can serve as resources for future studies for crop engineering and elucidates greater evolutionary dynamics of the extended NLRs than previously assumed.

## Introduction

Plants have evolved elaborate innate immune systems against invading pathogens[1]. As one of the major classes in plant immunity, resistance genes (R genes) encoding nucleotide-binding and leucine-rich repeat receptors (NLRs) induce a programmed cell death called hypersensitive response (HR) and downstream resistance by direct or indirect recognition of the effectors from pathogens. Plant NLRs in general, in contrast to animals’, evolved to form large gene families with hundreds of members, which display various multi-domain architectures (MDAs)[2, 3]. The conserved components of the MDAs are a nucleotide-binding domain shared by Apaf-1, resistance proteins and CED4 (NB-ARC or NB) and a variable number of C-terminal leucine-rich repeats (LRRs). The N-terminal domains are diverse to include a Toll/interleukin-1 receptor homology (TIR) domain, a Coiled-coil (CC), or a Resistance to Powdery Mildew 8 (RPW8) domain. Thus, the N-terminal domain typically determines the class of NLRs as TNLs (TIR-NB-LRRs) and non-TNLs which are further categorized as CNLs (CC-NB-LRRs) and RNLs (RPW8-NB-LRRs). Recently, additional integrated domains (IDs) which may function as baits for effector recognition have been identified in NLRs of flowering plants[4, 5]. In Solanaceae, a novel domain named Solanaceae Domain (SD) was found to be integrated at the N-terminus of CNLs and function for oligomerization or interaction with a decoy or effectors from pathogens[6–8].

Recent studies in genome-wide comparative analyses of NLRs provide insight for the dynamics of NLR evolution[9, 10]. Challenged by fast-evolving pathogen effectors, NLRs evolved to increase the copy number by tandem and segmental duplication and enhance genetic diversity by polyploidization and natural selection which sometimes also led to gene losses[11–13]. Significant involvement of retroduplication was demonstrated through the study of the NLRs in pepper in Solanaceae family, which showed that enormous clade-specific duplications occurred independently from other closely related species [14, 15]. A new paradigm of functional evolution of NLRs also recently appeared[16, 17]. It is that singleton NLRs evolved through specialization and diversification to be “sensors” detecting activities of pathogens and “helpers” executing the HR, which has eventually formed an extensive a helper-sensor network. In Solanaceae, multiple NLRs in the same clade to which NLR-required for Cell Death (NRC) protein belongs are known to function as helpers in coordination with NLRs from some other clades postulated to function as sensors[16].

Tomato (*Solanum lycopersicum*) in Solanaceae family is one of the most important vegetables in the world and challenged by over 200 diseases caused by diverse pathogens that can lead to great economic losses[18]. Owing to low genetic diversity and changes of resistance genes of the cultivated tomato during domestication, the necessity for genetic improvement has escalated[19, 20]. Hereditary resistance of wild tomato species against diverse diseases has obtained particular attention for this purpose[19, 21]. To date, tomato has 12 closest wild relative species[22], among which *Solanum pimpinellifolium* and a few other species have been known as the richest sources for disease resistance genes[19, 23]. Over the past decades, the perpetual effort has led to the identification of functional resistance genes, including NLRs, from wild tomato species and their successful introgression to the domesticated tomato[24]. As secureness and pyramiding of R genes were suggested for more durable resistance against multiple pathogens[25, 26], the value of the NLRome has been enhanced to search for functional R genes already developed in diverse habitats, from volcanic islands to over 3000m above sea level in The Andean Mountains[27]. Although there was a populational study of *Solanum chilense*[28], studies are still limited and no proper resources are yet available for high-quality NLRs owing to the financial and computational burdens.

Resistance gene enrichment sequencing (RenSeq) was recently developed as an efficient, cost-effective method to selectively capture and sequence NLRs in the species of interest[29, 30]. However, limitations existed as short reads from the Illumina platforms could not completely resolve highly repetitive sequences and physical clusters of NLRs[31]. Combined with single-molecule real time (SMRT) sequencing of PacBio, RenSeq has enabled more accurate recovery of NLRs and been successfully applied for the discovery of new functional R genes and evolutionary study of diverse accessions of *Arabidopsis* species[32]. In this study, we employed SMRT RenSeq to identify NLRs from 18 Solanaceae accessions (*S. lycopersicum* Heinz, 15 wild tomato accessions belonging to 5 species, *Nicotiana benthamiana* and *Capsicum annuum* ECW20R). We suggest the origin of the CNLs with extended N-terminal domains called SDs predates the divergence of Solanaceae and this domain is likely more structurally and functionally diverse. In addition, we demonstrate that the extended CNLs are a major component of NLR evolution in tomatoes, resulting in high variations in the number between wild tomatoes and displaying evolutionary signatures from interaction with pathogens. These findings may enhance our understanding of dynamic evolution of NLRs and provide the novel basis for disease resistance for crop breeding in Solanaceae.

## Results

### RenSeq for exploration of NLRs in Solanaceae

To identify NLR repositories and elucidate their evolutionary history in tomato, we utilized SMRT RenSeq to selectively sequence and annotate the NLRs of 15 accessions from five self-compatible wild species: *Solanum galapagense, Solanum cheesmaniae, S. pimpinellifolium, Solanum chmielewskii and Solanum neorickii*, and a domesticated tomato, *S. lycopersicum* Heinz (Supplementary Figure 1). For the comparison between genera in Solanaceae, NLRs of *C. annuum* ECW20R and *N. benthamiana* were generated through the same pipeline (Supplementary Table 1). SMRT RenSeq using a PacBio platform made it feasible to overcome the limitations of Illumina RenSeq, enabling accurate recovery of NLRs and their clusters (Supplementary Table 2). Each gene model was manually curated to enhance the quality of annotation. Combined together, our pipeline produced 264 to 332 high-quality NLR gene models in tomato, providing an important resource for future studies (Supplementary Table 3). In comparison to the reference genome of *S. lycopersicum* Heinz, we annotated 314 NLRs, among which 276 NLRs were almost identically identified in the annotation of ITAG3.2. Our RenSeq results improved the annotation of 128 NLRs including 13 existing annotations which were incomplete because of mis-assembly or unfilled gaps (Supplementary Figure 2). We only found three NLRs absent in our annotation set. Two of them were most likely pseudogenes short in length, and the other contains NB-ARC domain with low homology (1E-05). The improvement was much more noticeable in *N. benthamiana* for which we annotated 307 NLRs. In comparison to the currently available data containing 199 and 171 NLRs, respectively[33, 34], not only were more NLRs annotated but also their MDAs and lengths were significantly improved (Supplementary Table 4).

The annotated NLRs of tomato species were initially classified into the four representative classes based on the predicted N-terminal domains or motifs (Main Figure 1a). CNLs were predominant, followed by NLs to which putative pseudogenes that have lost their original MDAs were more frequently included (Main Figure 1b). TNLs accounted for about ten percent of the total number of NLRs. Only two RNLs were consistently annotated as N requirement gene 1 (NRG1) and Activated Disease Resistance 1 (ADR1) orthologs. Intraspecies and interspecies variations in the number of NLRs existed. *S. galapagense* and *S. cheesmaniae*, endemic to the Galapagos Islands, tended to encode a smaller number of NLRs than did the other three species in the continent of South America (Supplementary Figure 1, Supplementary Table 4). *S. neorickii* LA1322 sharing the habitat with *S. chmielewskii* in Southern Peru encodes more NLRs than did the other two accessions in Northern Peru and Southern Ecuador. Despite possible correlations of the expansion and contraction of NLRs with habitats or lineages, no statistical inferences were drawn due to the insufficient number of samples.

**Main Figure 1.**
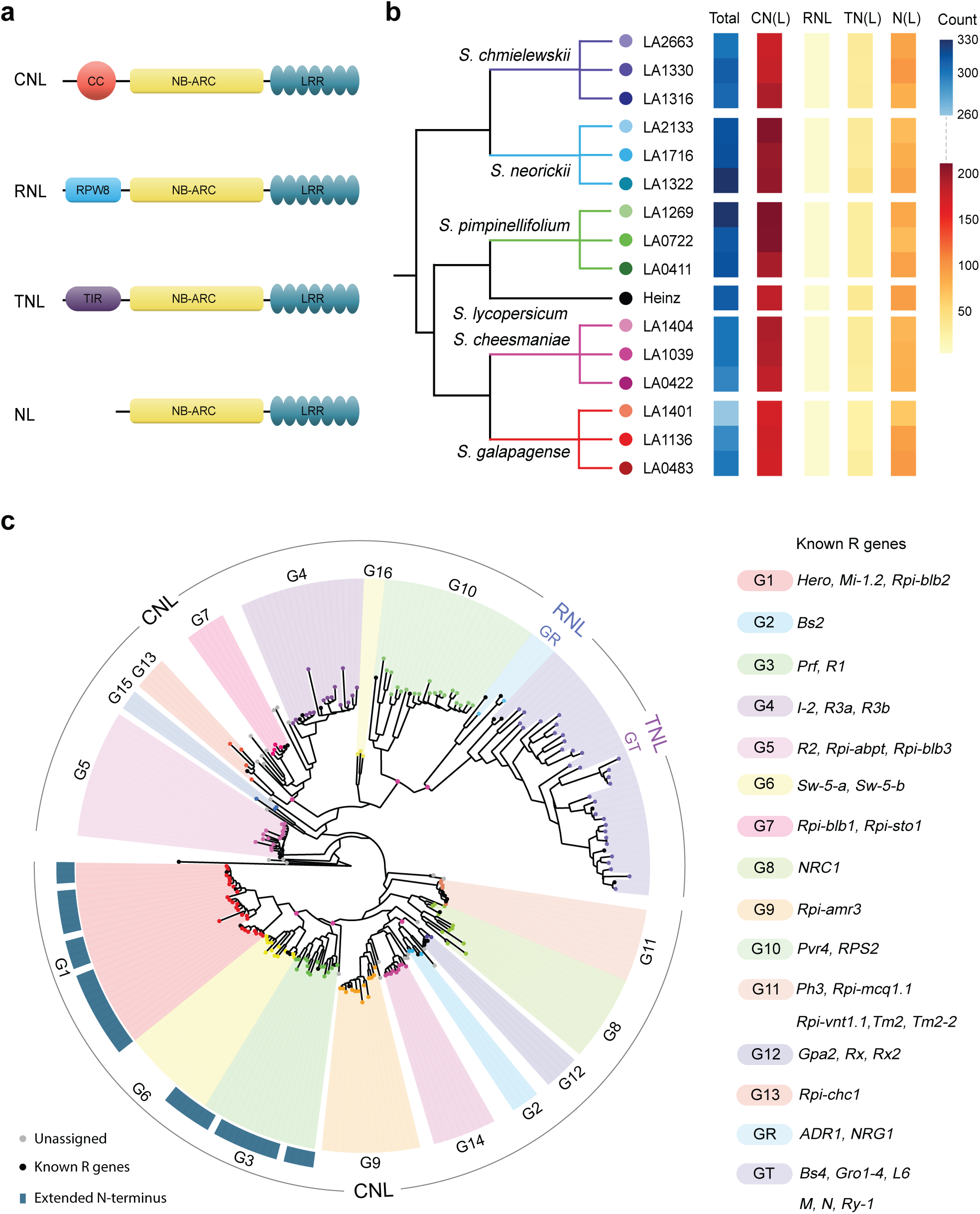
The diversity of NLRs in tomato. **a.** Schematics of the four representative multi-domain architectures of NLRs **b.** A putative species tree of the tomato species based on the previous study[100] and the number of NLRs in each representative class shown as a heatmap. **c.** The phylogenetic tree inferred using intact NLRs of S. lycopersicum Heinz. Clades were assigned based on bootstrap values (>=75) guided by the known R genes. Any nodes under which two or more clades existed with a significant statistical support (>= 75) were indicated with pink circles. NLRs having extended N-terminus were indicated with dark green rectangle. The representative classes of NLRs were given in the outer ring.

The NLRs were also classified based on their phylogenetic relationship. The NB-ARC domains containing at least three conserved major motifs among P-loop, GLPL, Kinase2 and MHD detected by NLR-Parser[35] over 160 amino acids were selected as intact NB-ARC and used for phylogenetic tree construction. The phylogenetic tree displayed a clear separation between the classes (Main Figure 1c, Supplementary Table 5). The TNL clade, GT, and the RNL clade, GR, which formed a monophyletic group were clearly distinguishable from the other CNL clades. Only the CNL clade, G10, consistently shared the most recent common ancestor (MRCA) with them, alluding to their evolutionary connection. The exact branch order of the other CNL clades could not be statistically confirmed in most cases due to the divergence of the NB-ARC domains. Nonetheless, the general topology agreed with the tree in the previous study in that G1 to G12 in the lower part of the phylogeny were clustered together with a helper clade (G8) to form a so-called NRC-dependent superclade (Main Figure 1c)[16]. Certain clades, however, were invariably clustered together with statistical significance. Among them were G1, G3 and G6 where NLRs with extended N-terminus were exclusively identified (Main Figure 1c). The N-termini of these NLRs were typically characterized with 550 or more amino acid sequences, about 350 amino acids longer than that of the typical CNLs. This pointed to the possibility that the extended N-terminus may accommodate additional structural or functional domain(s) other than the canonical domains of NLRs, which according to the phylogeny might have originated from a single ancestral NLR.

### Origin and diversity of the extended CNLs

To elucidate the diversity and evolution of the extended CNLs, the NLRs in CNL-G1, G3, and G6 were used for the detailed analysis. These three clades include several cloned functional R genes, namely Hero, Mi-1.2 and Rpi-blb2 in G1, Prf and R1 in G3 and Sw-5 in G6 (Main Figure 1c)[14]. The terminology for the extended N-terminus first appeared as Solanaceae Domain (SD) based on homology of Prf to these known R genes in Solanaceae[6]. A previous study followed this notion and reported a possible common origin of the NLRs with SD in pepper, potato and tomato in Solanaceae[14]. However, the pairwise similarity search between Hero or Sw-5 and Prf in fact confirmed homology only over CC-NB-LRR but not the N-terminus. To detect their distant homology by a more sensitive method, we generated a Hidden Markov Model (HMM) for each clade using manually curated extended N-terminal regions from our RenSeq data and searched for each on the other two clades[36]. These results supported strong homology between the extended N-terminal regions of G1 and G6 as each HMM was able to detect its homologous domain in nearly all the N-terminal sequences of the other clade (Main Figure 2a). However, homology between G3 and G6 was absent, and only the HMM of G3 was able to detect homologs in G1. This homology only occurred partially over a region of 200 amino acids, pointing to significant diversification on the other regions. Homology of G3 and G6 to the intermediate, G1, was the evidence for the distant homology between G3 and G6, suggesting a common origin of the N-terminal sequences of the three clades. However, that the NLRs in G3 would accommodate a globally homologous domain at the extended region as those in G1 and G6 was questionable.

**Main Figure 2.**
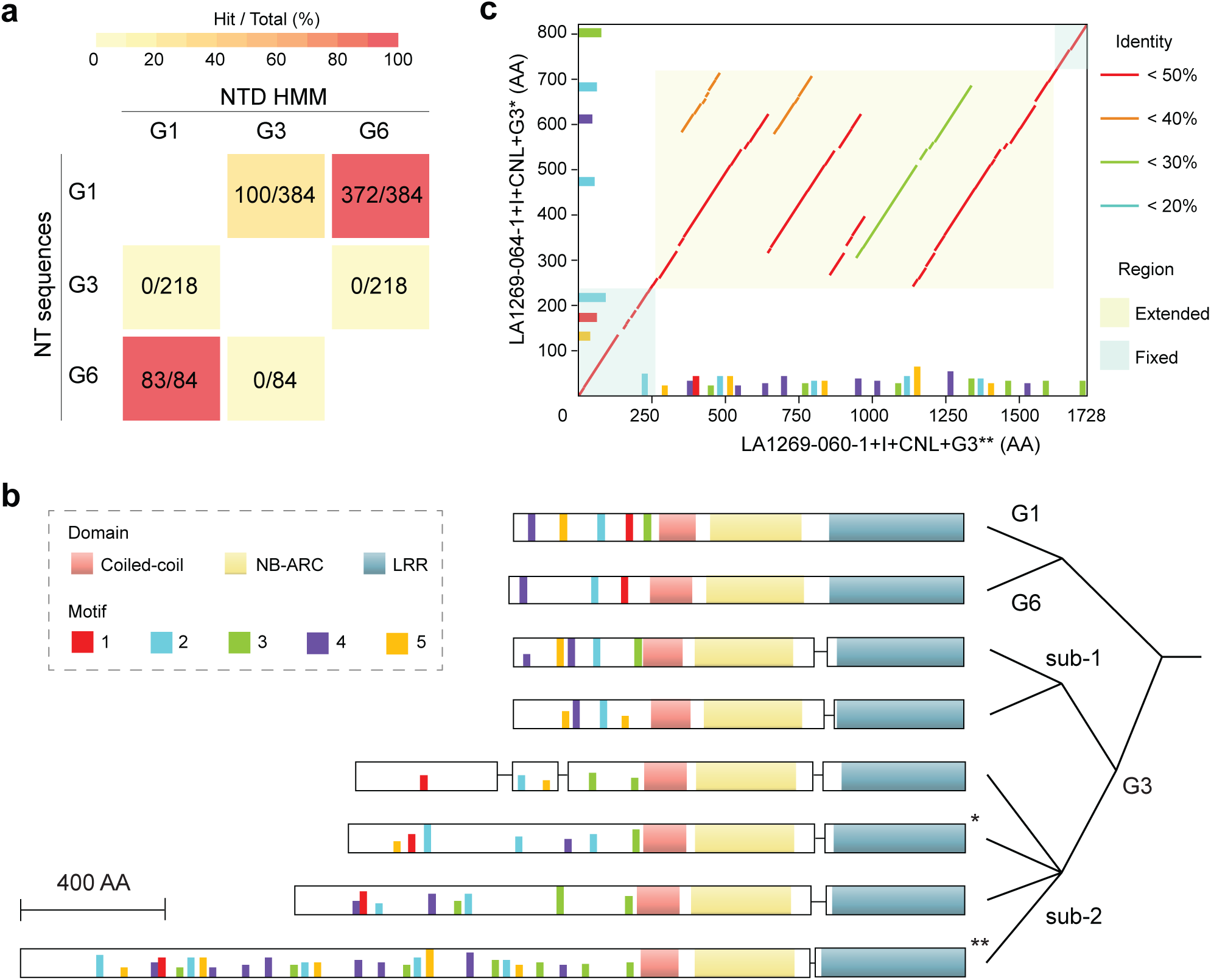
Sequence and motif analysis of the extended N-terminal domains. **a.** HMM search of N-terminal sequences. HMMs were built for the N-terminal domain (NTD) of each clade and searched on the other clades. The number of N-terminal (NT) sequences and the number of matches were indicated and visualized as a heatmap. **b.** Motif analysis of NLRs having extended N-terminal sequences. Five conserved motifs in the N-terminal regions (Supplementary Figure 3) and domains were depicted on the representative NLRs in G1, G3, and G6. Exons and introns were indicated as rectangles and lines, respectively. The gene structures for NLRs in G1 and G6 were highly variable. **c.** N-terminal sequence comparison between two NLRs in G3 subgroup-2. The location of the motifs and the sequence identity of the homologous regions were indicated in the dot plot.

We further performed motif analysis using Multiple Em for Motif Elicitation (MEME)[37] to understand the process of diversification of G3. There were five highly conserved motifs shared on the N-terminus by G1, G3 and G6 (Supplementary Figure 3). NLRs in G1 and G6 could be represented with a single set of motifs for each, resulting from a low degree of diversification of the motif regions within the clade (Main Figure 2b). In case of G3, the sequences formed two subgroups displaying different evolutionary patterns (Main Figure 2b). The N-terminal sequences in subgroup-2 of G3 were longer than those of typical extended CNLs in G1, G6 and even subgroup-1 of G3. This subgroup contained gene models of 1600 to 2600 amino acids characterized with a substantial difference in length for NLRs. The distribution of motifs suggested possible duplications of a certain set, contributing to the increase in the size of the proteins. This was clearly indicated in the comparison between the largest and the second smallest proteins in this subgroup, namely LA1269-060-1+I+CNL+G3 (2618 amino acids) and LA1269-064-1+I+CNL+G3 (1687 amino acids) (Main Figure 2c). The pairwise homology search result of their extended N-terminus identified in the beginning and at the end the “fixed” regions in which the homology remained almost intact. Between them was an “extended” region in which proliferation of the motifs resulted in repeated homology between the two proteins. The multiple duplications of the motifs into this extended region were regarded as the mechanism for the subsequent extension of the N-terminus without interrupting the original exon-intron structures (Main Figure 2b). Based on the homology between the motifs with low sequence identities in the extended N-terminus of LA1269-060-1+I+CNL+G3, it is thought that the diversification of the newly emerged motifs was concomitant (Supplementary Figure 4). On the other hand, subgroup-1 in G3 displayed a limited degree of elongation, only presenting gene models of 1200 amino acids, typical for the extended CNLs in G1 and G6, to below 1500 amino acids with the distribution of motifs highly resembling that of G1.

To trace the origin of the domains, we identified putative NLRs with extended N-terminus (300 or more amino acids prior to NB-ARC domain) in the selected dicots and scanned their N-terminus with the previously constructed HMMs. This analysis revealed the presence of homologous domains outside Solanaceae (Main Figure 3, Supplementary Table 6). About 80% of the extended N-terminus of NLRs in coffee in Rubiaceae were detected to have homology to the N-terminal domains in G3 (E-value < 1E-10). Olive and common ash in Oleaceae and sugar beet in Amaranthaceae also encoded a few NLRs with homology to the N-terminal domains in G3. Especially in sugar beet, the two matches displayed significant homology over 200 amino acids. As homologs were absent in Rosids, such as cassava, strawberry and *Arabidopsis thaliana*, the emergence of an ancestral domain was estimated to the MRCA of Asterids and Amaranthaceae (Main Figure 3). The common origin of these extended CNLs were further supported by a phylogenetic tree inferred with their NB-ARC domains (Supplementary Figure 5). The NLRs outside Solanaceae with their N-terminal domains sharing homology with those of NLRs in G3 formed a supercluster with the extended CNLs in G1, G3 and G6 in Solanaceae with a bootstrapping value of 92. The extended N-terminus in NLRs in coffee showed the best homology to both subgroup-1 and subgroup-2 of G3. However, outside Rubiaceae, partial homology to subgroup-2 was predominantly observed, possibly suggesting that this subgroup has most retained the ancestral sequence characteristics. We speculate that as suggested in the phylogenetic tree (Main Figure 1c), some ancestral NLR in G3 became fixed and proliferated to form new clades, G1 and G6, in Solanaceae and other independent or significantly diversified clades in the species outside Solanaceae.

**Main Figure 3.**
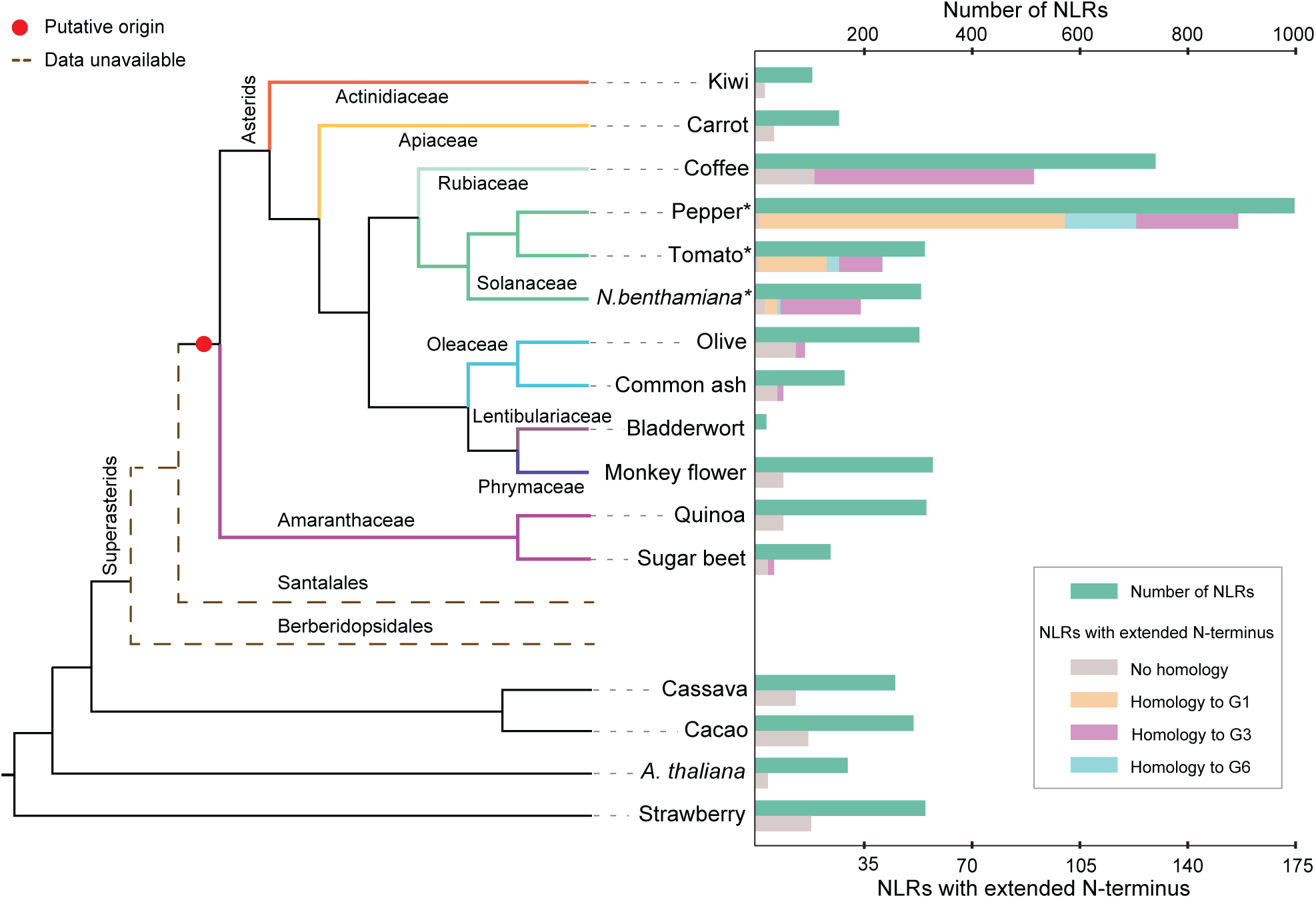
The origin of extended N-terminal domain. The numbers of NLRs and NLRs with extended N-terminus were indicated on the cladogram inferred using OrthoFinder and STAG[38]. The HMMs constructed for the N-terminal domains of the three clades, G1, G3 and G6, were searched on the extended NLRs. The best matches (E-value <= 1E-10) were counted and indicated. Hypothetical branches were added for Santalales and Berberidopsidales of Superasterids as their genome data was yet unavailable. The probable origin of the domain was indicated with a red dot. *Our RenSeq data was used instead of the genome data for *C. annuum* ECW20R, *S. lycopersicum* Heinz and *N. benthamiana*.

### Evolution of the extended CNLs in wild species tomatoes

To conduct in depth evolutionary study, we manually curated orthogroups (OGs) in 16 *Solanum* species guided by OrthoFinder[38] and phylogenetic trees inferred for each clade by RAxML (Supplementary Table 7). 413 OGs were identified in total. OGs with 16 members were the most abundant as 138, among which 134 were single-copy OGs (Supplementary Figure 6a). Only 18 OGs occurred as singletons. 49 OGs (11.9%) were assigned for G1, only 5 of which were single-copy OGs (Supplementary Figure 6b). This proportion was the lowest among the clades with more than 10 OGs assigned, suggesting a high degree of variation between the species and accessions. 296 OGs (71.7%) included NLRs from the domesticated species, *S. lycopersicum* Heinz, which were utilized to anchor those OGs to the putative loci in the reference genome. Other OGs were anchored, if possible, when the OG included a member from a multi-NLRs contig and at least one of the other members was already anchored into the genome. 342 OGs (82.8%) were in total anchored to the reference genome (Supplementary Figure 7). Chromosome 4 contained the greatest number of OGs as 77 while chromosome 3 only had nine. Overall, OGs that displayed variations in the number of NLRs between or within the species were anchored in chromosome 4, 5, 6, and 9. There was no correlation between unanchored OGs and their clades, but they seemed to be attributed to the evolutionary distance as those OGs tended to be predominant with NLRs from a specific set of species, for instance, *S. neorickii* and *S. chmielewskii*. We primarily focused on 248 OGs (60%) in which 80% or more members contained NLRs predicted to include intact NB-ARC domains. This was to avoid impacts from putative pseudogenes characterized with the anomaly of their NB-ARC domains, an unusual codon usage based on the principal component analysis (PCA) (Supplementary Figure 8a), and a potential loss of the original MDAs (Supplementary Figure 8b), which were all likely attributed to the accumulation of frequent random mutations[39].

A significant number of non single-copy OGs still remained after the filtering and represented a major portion in some clades including G1, G4, G5 and G6 (Supplementary Figure 6b). In PCA with all the non single-copy OGs, intraspecies clusters were formed with clear interspecies separations, except for the two closely related species in the Galapagos Islands, *S. galapagense* and *S. cheesmaniae* (Supplementary Figure 9). This suggested that the changes within the OGs were mostly correlated to major speciation events. To examine the possible source of the intraspecies variations, we obtained the genomic sequences of 48 additional accessions of *S. pimpinellifolium* and mapped them into the contigs of *S. pimpinellifolium* LA1269 (Supplementary Figure 10a, Supplementary Table 8) [20,40–42]. The presence and absence variations (PAVs) were inferred with SGSGeneLoss[43] and used for PCA. The accessions collected from Peru spread out without a clear correlation with geography, but those obtained from the northern Ecuador were more distinctly separated from the others (Supplementary Figure 10b). This tendency was in consistency with the previous observation[20] and suggested that intraspecies variations were the outcomes of the continuous local adaptation and/or population migration. These evolutionary and adaptive processes resulted in the greatest variation in the number of NLRs in G1 with 35 NLRs in *S. pimpinellifolium* LA0722 and LA1269 as the highest and 17 NLRs in *S. chmielewskii* LA2663 as the lowest (Supplementary Figure 11, Supplementary Table 9). This was followed by G4 with *S. neorickii* displaying a larger repository, GT with noticeable deviations of a few accessions and G6, whereas most of the other clades did not display significant differences.

The greater number of NLRs in *S. pimpinellifolium* was partially correlated with more gene copies in OGs but also attributed to the flexibility of its evolution. We found that OGs with duplicated NLRs from or exclusive to *S. pimpinellifolium* were rare (Supplementary Table 7). This suggests that ancestral NLRs in the OGs were present in some other species but partially lost in evolution. NLRs of *S. pimpinellifolium* also displayed flexibility in clustering. They tended to cluster with those of *S. galapagense* and *S. cheesmaniae* as in OG024, which are evolutionarily more closely related (Main Figure 1b, Main Figure 4a). But at the same time, as in OG030, some OGs were exclusive to *S. pimpinellifolium*, *S. neorickii* and *S. chmielewskii* in the continent of South America, collectively alluding to both evolutionary and geographic pressures (Supplementary Figure 1, Main Figure 4a). We explored genomic organization of NLRs in G1 in depth. Most of the OGs of G1 were anchored in chromosome 4, 5 and 6 (Main Figure 4a). OGs anchored in chromosome 4 were consistently clustered together in the phylogenetic tree, suggesting translocation of these OGs between chromosome was unlikely. However, the other two subclades displayed clustering of OGs anchored in different chromosomes. Interestingly, the OGs in chromosome 4, nevertheless, displayed the most significant variations (Main Figure 4b). A gene cluster spanning 83 kb included an OG from the NRC clade known as helper (G8) in the center and multiple OGs of G1 tandemly located in a complex manner. This region seemed to have undergone massive duplication and/or deletion events. No correlation between phylogenetic relationship and physical distances were found, and the number of NLRs and OG compositions by accessions also varied significantly. Transposable elements (TEs) such as LTR retrotransposons copia-type found in this region supported that these NLRs were not simple outcomes of tandem duplication but the transposon activities could have been involved in the birth, death and rearrangement of NLRs perhaps in lineage-specific manners. Some NLRs displayed single-exon gene structures, likely outcomes of retroduplications. But the gene structures were not necessarily consistent within the OGs, interrupted by mutations altering the exon-intron junctions. Nevertheless, NLRs of G8 in this cluster were conserved among all species, possibly suggesting that NLRs in G1 purposely expanded around the helper NLR.

**Main Figure 4.**
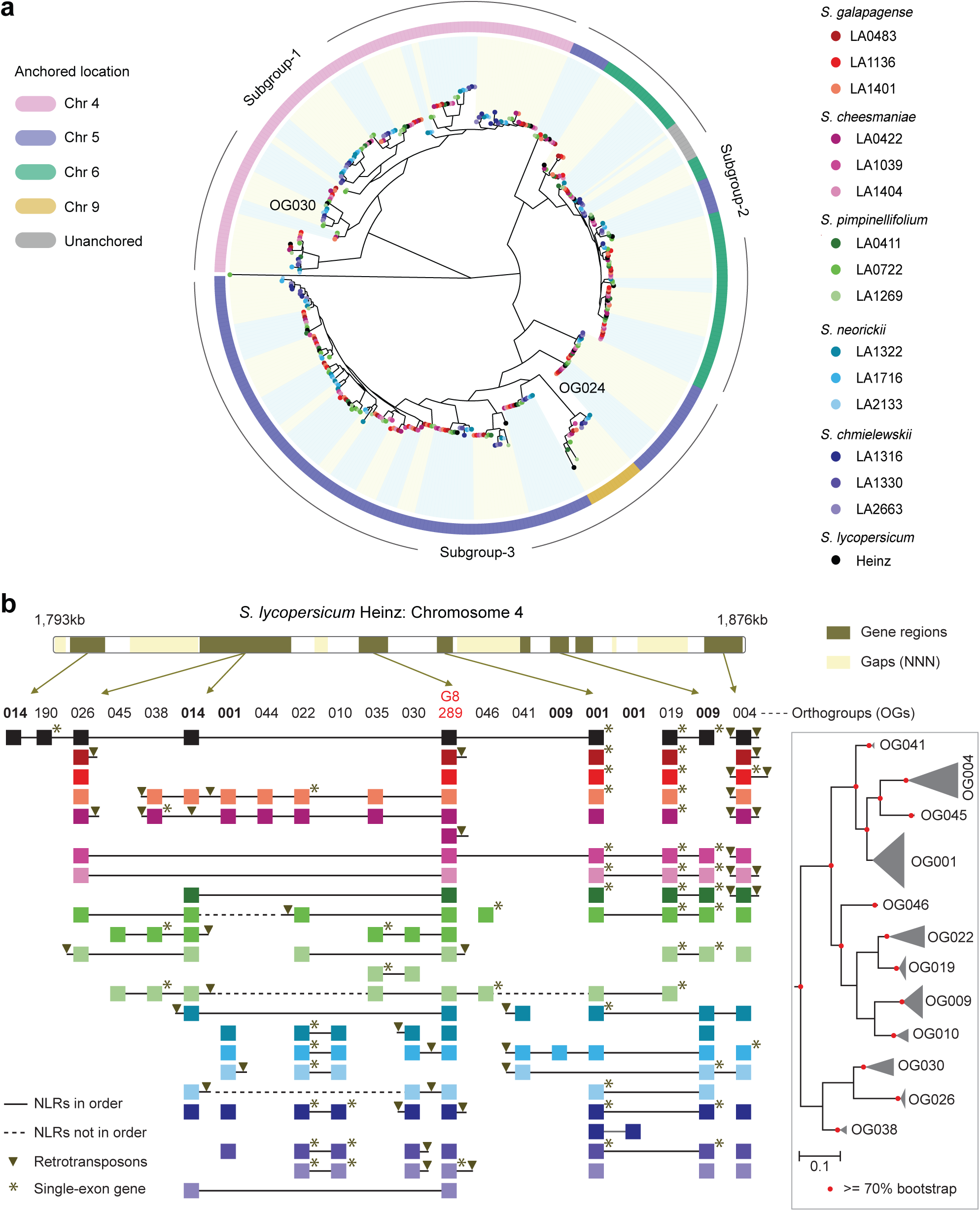
Diversity of NLRs in G1 clade. **a.** A maximum-likelihood tree inferred by RAxML using the NB-ARC domains of NLRs from selected OGs that belong to G1 (See results for selected OGs). The tree was rooted with an outgroup. The OGs were indicated using alternative background colors. The OGs anchored to reference genome were indicated in the outer ring. **b.** A representative example G1 physical cluster in chromosome 4. The OGs and their members anchored to this region were depicted. OGs except OG289 (G8, intact) and OG190 (G3, partial) are in G1 clade. NLRs in the same contig were connected by solid (in order) or dashed (not in order) lines. Retrotransposons near the NLRs were indicated as an inverted triangle. NLRs having a single-exon gene structure was indicated with asterisk (*). A relevant portion of the phylogenetic tree was given. Nodes with a significant statistical support (>=70) were indicated with pink circles.

### Diversification and selection in the extended CNLs

Such high variations in G1 may be attributed to the recent lineage-specific duplications. We estimated the pairwise synonymous substitution rates (Ks) for each pair of NLRs in the same clade for each species. As the amino acids significant for function and structure are often under negative selection, pairwise Ks of paralogs would typically continue to increase until saturation in the course of evolution. The comprehensive Ks plot composed of representative Ks revealed that their majority in G1 fell below 0.2, and Ks below 0.05 were highly enriched in comparison to the other clades (Supplementary Figure 12). This pattern indicated that the expansion of G1 likely began sometime after the speciation event from the MRCA of tomato and pepper, estimated as Ks of 0.3[44], and the variation in the number of NLRs was attributed to more recent lineage-specific evolution in tomato. G4, G5, and G6 indicated to show variations between the species also displayed greater number of recent duplication events than that of the other clades that showed no noticeable variation. Despite the relatively recent emergence of the OGs in G1, the global Ks and non-synonymous substitution rates (Ka) estimated by CodeML[45] suggested they could be under high positive selections (Supplementary Figure 13). Overall, the Ka/Ks of most OGs remained below 1, as the increase in Ka and Ks was accompanied. However, the increasing Ka pointed to greater adaptational signals. Although this was the most noticeable in the C-terminus, the N-terminus, including the extended domains and CC, and the NB-ARC domain were also subject to be under positive selection pressures.

We combined the information of the estimated time of duplication events and the divergence within OGs to illustrate their evolutionary pattern (Main Figure 5a). Each OG with six or more members were plotted based on the average pairwise Ks of the members to the closest outside the OG and Shannon’s entropy derived from protein sequence alignments. Despite an ancient duplication event, the two OGs in GR, which are orthologous to NRG1 and ADR1, evolved to maintain a substantial level of sequence conservation within the OGs. Then, OGs that belong to the clades in the upper part of the phylogeny, including G4, G10 and GT, began to appear, suggesting that they did not undergo duplication events after a relatively ancient one (Main Figure 1c). Between the average Ks of 3.5 to almost 1, OGs in GT were predominantly observed, affected by yet unknown pressures for clade expansion. Many of these OGs displayed a relatively low level of entropy despite their more ancient emergence, indicating that the sequences were generally conserved. Similarly, OGs in G8, including the one orthologous to NRC1, between the average Ks value of 2 and 0.5, indicated limited recent expansion and high sequence conservation. OGs in G3, including the one orthologous to Prf, started to increasingly appear past the average Ks of 1. Other extended CNLs in G6 and G1, on the other hand, were characterized with much smaller average Ks because of recent expansion of the clades. The average Ks below 0.5 was characterized by exponential duplication events of CNLs clades. Although the most recent duplications were predominantly represented with OGs in G1, other clades, such as G4 and G5, underwent significant expansion. In accordance with the previous observation, the increase in entropy indicated that the sequences within these OGs were much more diverse, alluding to evolutionary pressure to result in such variations.

**Main Figure 5.**
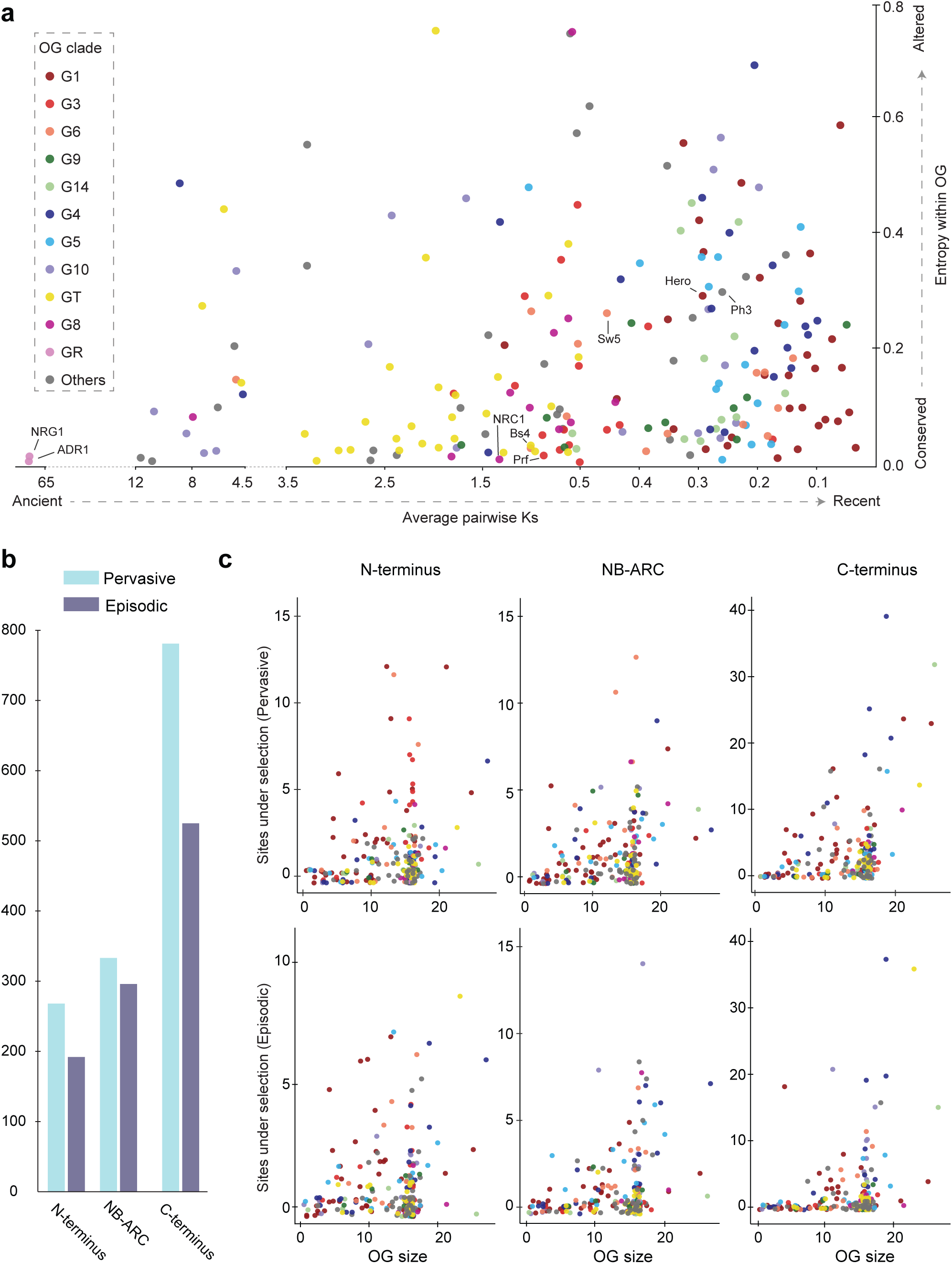
Diversification and positive selection pressures of the OGs. _a._ Estimation of the relative time of duplication events and sequence diversification within OGs. Each colored dot is an OG categorized based on its clade. The extended CNLs (G1, G3 and G6), CNL clades in the lower half of phylogeny (G9 and G14), CNL clades in the upper half of phylogeny (G4, G5 and G10) and helper clades (G8 and GR) were given similar colors, respectively. The x-axis indicates the average pairwise Ks values of the members in an OG to the closest outside the OG. The greater this value becomes, the more anciently the OGs were involved in duplication events. The y-axis indicates Shannon’s entropy within an OG derived from its protein sequence alignment. Entropy of zero refers to a perfect sequence conservation, and the increase in entropy means increase in sequence divergence. **b.** For the selected OGs, site-level pervasive and episodic selection pressures were detected in the N-terminus, NB-ARC domain and the C-terminus by FUBAR and MEME, respectively. **c.** The number of sites under pervasive (top) and episodic (bottom) selection pressures were categorized by OG size and clade.

The divergence within OGs could be correlated with statistically significant site-specific pervasive and episodic positive selection pressures using Hypothesis Testing Using Phylogenies (HyPhy)[46]. Overall, 781 and 525 sites in the C-termini were under significant pervasive and episodic selection pressures, respectively, indicating that LRRs have undergone both continuous and transient changes in response to environmental and evolutionary pressures (Main Figure 5b). Sequences in the N-termini and NB-ARC domain were, at a smaller degree, subject to the positive selection pressures. The OGs of extended CNLs in G1, G3 and G6 tended to display a greater number of sites under pervasive selection pressure possibly as the N-terminal domains may continuously evolve in response to pathogens (Main Figure 5c). The length of the extended domains was, however, not necessarily proportional to the number of sites detected. The OG with the longest gene model (2618 amino acids), for instance, only showed four sites under the pressure. Other N-terminal domains such as RPW8 and TIR were not under comparable level of selection pressures. The sites under pervasive and episodic selection pressures at the C-terminal LRRs showed high variations. Noticeably, some OGs displayed a significantly greater sites under the selection pressures as observed for the NLRs of *Arabidopsis*[32]. The clades in the upper half of the phylogeny, such as G4 and G10 (Main Figure 1c), appeared more frequently to have 10 or more sites detected. Nevertheless, 38% and 58% of the OGs were not detected for any pervasive and episodic selection pressures, respectively, leading to a curious question about how this trend would be related to evolutionary time and/or pressure.

## Discussion

In response to an increased risk of infection in domesticated tomatoes resulting from low genetic diversity and alteration of resistance genes, researchers have endeavored to identify functional R genes from wild tomato species already developed through co-evolution with their pathogens in highly diverse habitats [19,20,24,47]. However, the large number of NLRs in genome as well as technical and financial obstacles for genome assembly and annotation have hindered the generation of the comparative resource. In this study, we demonstrated that SMRT RenSeq is a cost-effective, efficient alternative to the whole genome sequencing. We also verified that SMRT RenSeq was capable of overcoming the limitations of Illumina RenSeq in resolving the complexity of NLRs and their clusters, resulting in a more complete set of annotations[30]. Our data showed significant improvement in comparison to the annotations of *S. lycopersicum* Heinz (ITAG3.2) and *N. benthamiana*, in support of the thoroughness of our pipeline and the reliability of our data (Supplementary Figure 2, Supplementary Table 4). We believe that manually curated annotations of NLRs for 18 accessions in Solanaceae will serve as comparative resources for the future studies.

We present an evolutionary study for the NLRs with extended N-termini identified in three clades, G1, G3 and G6 that would accommodate yet non-canonical domains of NLRs (Main Figure 1c). Although this putative domain has been commonly indicated as Solanaceae Domain (SD)[6,8,14], we consider this terminology could be a misnomer mainly for several reasons: the absence of a globally homologous domain between G3 and G1 or G6 and the emergence of the ancestral domain in the MRCA of Asterids and Amaranthaceae. We demonstrated through the disagreement in the organization of the motifs and the level of their sequence divergence between clades (Main Figure 2) that although the three clades originated from a common ancestral extended CNL, the extended N-terminal domain significantly diversified by different ways of selection and evolution. Our data suggested that the ancestral extended CNL seemed to start significant expansion before the divergence of coffee, and G3 was likely the initial outcome of that expansion (Main Figure 3, Supplementary Figure 5). Interestingly, the two homologs from sugar beet, Bv5_105980 and Bv5_105960, together with a member of the NRC clade (G8), Bv5_105990, were previously associated as a possible representative of an ancestral helper-sensor NLR pair unique to Asterids and Caryophyllales, the order that includes Amaranthaceae[16]. Their analysis, in concurrence with our interpretation, led to the conclusion of an exponential expansion of the NRC-dependent superclades prior to the speciation of Gentianales, which includes coffee, about 100 million years ago (Mya). Our analysis provides an additional context that one of the ancestral forms of sensor NLRs already contained an extended N-terminal domain, and the expansion of the extended CNLs contributed to that of the NRC-dependent superclades.

In tomatoes, three major modes of evolution exist for the extended CNLs: continuous elongation of N-terminal regions as in G3, rapid proliferation of NLRs and sequence diversification as in G1 and domain loss as in G6. We demonstrated that the gene models in subgroup-2 of G3 were a result of continuous elongation of the N-terminus, eventually resulting in a gene model with over 2600 amino acids, three times larger than typical CNLs (Main Figure 2b). The mechanism of elongation is thought to be similar to how LRR increases its numbers within the MDA by duplicating its segment and inserting it adjacent to the origin (Main Figure 2c)[48]. Such mechanisms were suggested to be complementary to low generation rates of plants so as to potentially offer advantages to evolve more interfaces for protein-protein interaction. Our postulation is that an ancestral copy underwent duplication event with one paralog fixed and the other eventually undergoing further elongation with the newly duplicated segments eventually becoming diversified (Supplementary Figure 4). An outcome of this process is Prf composed of 1824 amino acids. Together with serine-threonine protein kinases named Pto or Pto-like proteins, Prf induces the HR in response to AvrPto secreted by *Pseudomonas syringae* pv. tomato[6,49,50]. The previous studies support that the extended N-terminal domain serves as an oligomerization interface for self-association and interaction with kinases to signal for the HR[51, 52]. Collectively, these studies support the idea that the elongation process could be a strategy to evolve a novel function to interact with other protein components.

It will be intriguing to further study the evolutionary benefits and costs of this process and its outcomes. NLRs indirectly recognizing pathogens were suggested to have unfavorable fitness cost as they could be susceptible to mutations on them or their interacting partners that can lead to auto-immunity[53]. Furthermore, although wild tomato species displayed a great level of sequence conservation for these extended NLRs in general, the longest gene models of *S. chmielewskii*, for instance, were commonly characterized with substantial increase in exon numbers affected by 700 nucleotide deletion in the extended N-terminal region, indicative of pseudogenes (data not shown). In pepper, more dramatically, no gene models in G3 with over 1500 amino acids were found whereas *N. benthamiana* and potato still encode them. The NLRs in subgroup-2 of G3 typically anchored in chromosome 5 in tomato seemed to be lost during the massive genome evolution of pepper.

NLRs in G1 and some in G6, on the other hand, were characterized with significant proliferation and sequence variation. We demonstrated that G1 contributed at the greatest degree to the variation in the number of NLRs among species and even accessions (Supplementary Figure 6, Supplementary Figure 7). *S. pimpinellifolium* in Peru tended to encode the greatest number of NLRs in G1, which seemed to be affected by both speciation events and adaptation and migration of population (Supplementary Figure 11, Supplementary Figure 12)[20]. The recent expansion of G1 (Supplementary Figure 12) resulted in physical clusters in which durable functional genes are present, including Mi-1.2 in chromosome 6 and Hero in chromosome 4 (Main Figure 4a), which confer resistance against root-knot nematodes, aphids, whiteflies and psyllids[54–56], and cyst nematodes, respectively[57, 58]. A physical cluster of G1 in chromosome 4 is a representative example of diversification of G1 (Main Figure 4). Unlike NLRs in G3 which have conserved putatively ancestral gene structures (Main Figure 2b), NLRs in G1 displayed diverse exon-intron structures, including a single-exon model, suggesting that duplications were mediated by not only tandem duplications but also retroduplications as in pepper even if the mainly targeted clade is different[15]. Despite the dynamical evolution of G1, one OG in G8 was well conserved in this cluster, perhaps serving as a core of sensor-helper clusters[17]. We further supported that many OGs in G1 would function as sensors based on higher positive selection pressure likely resulting from evolution against rapidly evolving effectors of pathogens in spite of the relatively recent emergence. However, it is worthwhile to note OGs in TNLs were characterized with relatively ancient duplication events and higher sequence conservation than that of OGs of potential sensor CNLs, although TNLs include sensors such as Roq1 in *N. benthamiana* which directly recognizes a conserved effector XopQ from *Xanthomonas*[59, 60]. This raises an interesting question whether a relatively small degree of polymorphism would be sufficient to handle the evolution of effectors or whether effectors recognized by TNLs would be generally conserved across pathogen species. It is also worth noting that the pattern for the expansion and the diversification of OGs outside NRC dependent clades, such as G4 and G5, was similar to the OGs in G1. It will be intriguing to answer why certain clades were preferred by evolution for expansion and diversification.

We also observed that that some NLRs in G6 have lost the extended N-terminal domain and CC in the course of evolution (Main Figure 1c). In consideration of their physical clusters in chromosomes, it seems an ancestral NL tandemly proliferated to increase the copy number. If the N-terminal domain and CC are essential for auto-inhibition of NLRs as found for Sw-5 in G6[8,61–63], it is yet unclear how this ancestral NL achieved it without the two domains and was eventually selected by evolution. Although some attempted to elucidate their functions, more diverse studies will be required[64].

Our study established a foundation to investigate NLRs in tomato, especially CNLs with extended N-terminal domains. However, some questions still remain to be resolved. It is unclear whether the extended domain might have originated by *de novo*, as postulated for RNLs[65, 66], or tandem duplication within the gene was already driving the elongation from an ancestral CNL[48]. It is curious how structurally the new large domains would fit the recently identified structure of ZAR1[67] and also how functionally they would differ from closely related CNLs. We also look for future studies for self-incompatible species, such as *Solanum habrochaites* and *Solanum peruvianum*, known as great resources for disease resistance stemming from substantial genetic diversity[19, 47]. Despite their advantages, they were absent in this study as most accessions maintained a high level of heterozygosity unable to be completely resolved in our pipeline. Although Mi-1.2 was initially identified from *S. peruvianum*[24], it is yet unclear whether similar evolutionary dynamics exists for the extended CNLs in these species. A larger scale of comparative study including these evolutionarily distant wild tomato species will provide more comprehensive insights on NLR evolution and expand the scope of NLRome for genome engineering and breeding of tomatoes.

## Materials and Methods

### Plant materials and library preparation

The seeds of wild tomatoes were obtained from Tomato Genetics Resource Center (TGRC) at UC Davis, CA, USA. Plants were grown in a walk-in chamber at 25°C. Genomic DNA of young leaf tissues were frozen by liquid nitrogen and extracted using DNeasy Plant Mini Kit (Qiagen, CA, USA). 10 μg of gDNA were fragmented with the Covaris sonicator (Covaris Inc., MA, USA) and size-selected with Sage-ELF (Sage Science, MA, USA) to get 2-5 kb fragments. The fragmented gDNA was used for the generation of libraries with the NEBNext Ultra DNA Library Prep Kit for Illumina (NEB, MA, USA). We used different combinations of primers as barcodes to pool two different libraries into one SMRT cell. To capture the NLR gene fragments, customized MYbaits kit (Arbor Biosciences, MI, USA) was used. The captured libraries were amplified using PCR (KAPA HiFi enzyme) and again sized-selected with Sage-ELF. The multiplexed libraries were sequenced using PacBio platform conducted at the Vincent J. Coates Genomics Sequencing Laboratory (GSL) at the University of California, Berkeley.

### Circular Consensus (CCS) assembly

We used SMRT Link (www.pacb.com) to generate CCS reads from the sequencing output. The CCS reads required at least three full-pass subreads from the insert and 90% of accuracy. All parameters were kept default, except for the maximum length of the CCS reads adjusted to 50,000. The CCS reads were demultiplexed using a provided script (https://github.com/weigelworld/pan-nlrome/tree/master/results/scripts). The demultiplexed CCS reads without proper primers were detected using BLASTN and removed (-evalue 1E-10 -perc_identity 99 -max_target_seqs 1)[68]. Seventy nucleotide sequences were trimmed off from both ends of the CCS reads to remove the adapter and barcode sequences using Cutadapt v 1.16+2.gaaef7dd[69]. Each set of the demultiplexed CCS reads was assembled using Canu v1.6[70]. For the parameter, the genome size of 3.0 Mb was estimated based on the approximate number and the average length of NLRs in the genome of *S. lycopersicum* Heinz[42]. The other parameters were determined based on the statistics of the number of mis-assembly and recovered NLRs in contigs assembled with different combinations of the parameters. Each different set of contigs was aligned to the chromosomes of *S. lycopersicum* Heinz (SL 3.0), and the alignments were evaluated by Quast v4.5 for mis-assembly and the number of recovered NLRs[71]. The following parameters that produced contigs with the lowest number of mis-assembly and the largest number of NLRs were used for the assembly: --pacbio-corrected, genomeSize=3.0 Mb, correctedErrorRate=0.025, minOverlapLength=350, trimReadsCoverage=1 and minReadLength=1200.

### Annotation of NLRs

We used MAKER[72] to predict the gene models of NLRs. For each accession, we independently ran MAKER three times with various *ab initio* gene prediction tools trained in different manners. 1) Augustus[73] and Snap[74] were selected as *ab initio* prediction tools. Augustus was trained with a non-redundant set of genes from ITAG 3.1 produced with cd-hit[75] by removing protein sequences that had 80% or more identical matches in the proteome (-c 0.8 -n 5). The genes without atypical splicing sites or in-frame stop codons were removed, and the remaining gene models with 1 kb flanking regions were used for the training. SNAP was trained only with putative NLRs extracted from ITAG 3.1 and their 1 Kb flanking regions. 2) Maker was run without any *ab initio* prediction tools. Maker identified protein coding genes based on the alignment of cDNA and protein sequences to contigs (cdna2genome = 1 and protein2genome = 1). The gene models generated in this process were used to train SNAP. Maker was re-run with the trained HMM of SNAP and without the cDNA and protein alignment (cdna2genome = 0 and protein2genome = 0). This process produced improved gene models, and SNAP was trained anew with them. Maker was finally re-run with this re-trained HMM to generate the second set of gene models. 3) Genemark-ES[76] was self-trained with the genomic sequences of contigs (--ES -min_contig 1). The trained HMM was used in MAKER to produce the last set of gene models. The protein sequences from the three sets of gene models were searched by InterProScan v5.25-64.0[77] to identify associated domains, namely TIR (PF01582 and PF13676), NB-ARC (PF00931), RPW8 (PF05659) and NB-LRR (PF12061) using the PFAM library v 31.0[78].

In addition to *ab initio* prediction, we used the CDS and protein databases as the evidence of the predicted gene models. The annotation sets included *S. lycopersicum* Heinz ITAG 3.2[42], *S. pennellii* v2.0[79], *S. tuberosum* Group Phureja v3.4[80] and *C. annum* CM334 v1.55[44] as well as the NLRs from the Plant Resistance Genes database (PRGdb) v2.0[81]. We added the annotation set of wild tomato species we generated into the database whenever the annotation became complete. This database was commonly used for all the runs of MAKER.

Repeat masking was conducted in MAKER using RepeatMakser v4.0.7 (http://www.repeatmasker.org). The miniature inverted-repeat transposable elements sequences of *S. lycopersicum* and *S. tuberosum* from P-MITE[82], transposable element protein sequences from MAKER (te_proteins.fa) and RepBase[83] were used as the database. Soft masking was enabled to avoid over-masking of NLR regions.

InterProScan was also run on the contigs. InterProScan translated the contigs in the six frames and identified all possible open reading frames (ORFs) equal to larger than 30 amino acids. The ORFs were searched against the PFAM to identify NB-ARC domain.

### Manual Annotation

The gene models from three runs of MAKER, evidence tracks and the predicted domains on the gene models as well as the translated contigs were visualized in Apollo genome browser v2.0.6[84]. We manually curated all the gene models that contain the NB-ARC domain detected by InterProScan or accession-specific HMMs of NB-ARC domains we designed. We followed the consensus of the predicted gene models. When there was no consensus in the gene models, we primarily followed cDNA alignment and other evidence. If no proper evidence was present, we aimed at capturing a gene model with a longer, continuous NB-ARC region. For putative pseudogenes with highly fragmented gene structures, we aimed to capture the entire multi-domain architecture possible even though supporting evidence was absent. After the manual curation, we examined gene models that may have false duplicates introduces during genome assembly. We first identified all possible pairs of annotated genes which shared 99% or more identity in their genomic sequences at least over 300 nucleotide or in their protein sequences using BLASTN and BLASTP, respectively. We visually inspected the alignments of contigs, genomic sequences, exon-intron structures, protein sequences and genomic contexts of each pair and excluded one of the pairs if there was evidence that one of them was a false duplicate. The refined annotation set was compared to the predicted gene models by TGFam-Finder v1.01[85] and finalized.

### Identification of multi-domain architectures and conserved motifs in NLRs

The annotated proteins were classified based on their multi-domain architecture by the previously constructed pipeline[14]. Briefly, we generated a HMM of NB-ARC domain with the high-confidence species-specific NLR gene models. TIR, RP8W, LRR and CC motifs were predicted by SMART, Pfam and COILS[78,86,87]. OrthoMCL clustering was used for the classification of CC motifs[88].

To study the extended N-terminal domains, we first obtained the putative domain regions from Mi1-2, Prf and Sw5-a comparted in the previous studies and manually identified them in NLRs of S. pimpinellifolium LA1269. Using homology, we extracted the extended regions from all *Solanum* species in this study. For each clade, redundant sequences were removed using cd-hit (-c 0.8), and the remaining sequences were aligned using MAFFT (--maxiterate 1000 –globalpair)[89]. The MSA was used to construct a HMM. The clade-specific HMMs were scanned to identify putative matches in other dicot species (Supplementary Table 6) (--domE 1E-10 -e 1E-10) or in tomatoes (--domE 1E-4 -e 1E-4). We further used MEME v5.0.5 to identify conserved motifs in the extended N-terminus of the NLRs in G1, G3 and G6 (-minw 5 -maxw 20 -nmotifs 20)[37]. We manually selected the motifs generally conserved in the three clades.

### Phylogenetic classification of NLRs

To classify NLRs by their phylogenetic relationship, the final annotation set for each accession was evaluated for the presence of major (P-loop, GLPL, Kinase2 and MHD) and minor (RNBS-A, RNBS-B, RNBS-C and RNBS-D) motifs in the NB-ARC domain by NLR-Parser v0.7[35]. The sequences from the first appearing major motif to the last in the correct order were considered as NB-ARC domain. Based on the previous study[14], we defined an intact NLR to have a NB-ARC domain equal to or larger than 160 amino acids as well as at least three major motifs without the requirement of minor motifs. We manually added in the NB-ARC domain of ADR1 orthologs undetected by NLR-Parser using the alignment of the NB-ARC domain of NRG1 orthologs.

We initially created phylogenetic tree using the intact NLRs of *S. lycopersicum* Heinz, representative known NLR genes (Main Figure 1c) and CED-4 in *Caenorhabditis elegans* as an outgroup. The protein sequences were aligned by MAFFT v7.313 (--maxiterate 1000 –globalpair)[89]. The MSA was imported to MEGA7 v7.0.26[90], visually examined for any misalignment and manually corrected. Any columns containing gaps in 90% or more positions were removed using TrimAl v1.4.rev22[91]. RAxML v8.2.12[92] was used to infer a phylogenetic tree based on the maximum likelihood method. The PROTGAMMAJTTF model, which was predicted to perform the best in this dataset, was chosen with random parsimony seeds of 12345 to identify the best tree starting from 100 randomly generated maximum parsimony trees (-m PROTGAMMAJTTF -p 12345 -# 100). The best tree was supported with 500 replications of bootstrapping (-m PROTGAMMAJTTF -p 12345 -b 12345 -# 500).

We assigned clades based on the previous study[14]. The highest node of the clade which was supported with a bootstrapping value of 75% or greater. In addition, we selected reference NLRs from *S. lycopersicum* Heinz considering their exon-intron structures, the length of their NB-ARC domains and the size of the clades. For the other accessions than *S. lycopersicum* Heinz, the intact NLRs of that accession, the reference NLRs from Heinz, the known NLR genes and CED-4 were all used to infer a phylogenetic tree. The clade was assigned based on the same criteria. To overcome the inconsistency of the classification based on phylogeny, we used Markov clustering. The intact NLRs of tomatoes or peppers were searched in all-versus-all BLASTP (-evalue 1e-4). The matches with the percent sequence identity equal to or greater than 90% were retained, and Markov clustering was applied with mcl with the sequence identity as weight (-I 2.0)[93].

NLRs without intact NB-ARC domains (partial NLRs) were assigned to putative phylogenetic clades based on the similarity search. They were searched against all intact NLRs in BLASTP (-evalue 1e-4). If NB-ARC domain of a partial NLR shared 50% or more sequence identity with its match, it was assigned to the phylogenetic clade of that match. Otherwise, no clade was assigned to it.

### Orthogroup generation and refinement and mapping

We identified orthogroups (OGs) of all the annotated NLRs in tomato using OrthoFinder v2.3.1[38]. OrthoFinder initially called DIAMOND v0.9.22.123 for the all-verses-all similarity search of the protein sequences (--e 0.001 --more-sensitive)[94]. The bit scores were normalized based on the length of the proteins in each pairwise species comparisons and Markov clustering was applied to the entire protein network with mcl (-I 1.5).

Some of the OGs generated by OrthoFinder presented the tendency of over and/or under-clustering and required manual refinement. We speculated that OrthoFinder generated higher scores between the orthologs of similar lengths during its normalization of bit scores. This would result in truncated forms of orthologs assigned into the same OG separated from other fully annotated orthologs. To complement this problem, we collected OGs that are likely correctly generated based on the two criteria. First, none of the species were supposed to have more than one homolog present. Second, the members were not supposed to have matches outside their OG that share a sequence identity over 96% in the NB-ARC domain based on BLAST. We collected all the other sequences that fail to meet these criteria to manually confirm them. Based on the previous annotations on phylogenetic clades, the protein sequences were separated. For each clade, a filtered MSA was generated for the entire MDAs of the proteins aligned using MAFFT and trimmed with TrimAl (-gt 0.2). A phylogenetic tree was inferred using RaxML (-p12345 -# 100 -m PROTGAMMAJTTF) with 250 replications of bootstrapping. We manually curated each OG using the statistical supports of the tree, pre-defined OGs by OrthoFinder and the sequence similarity between the members. Each curated OG was aligned with prank[95] and the MSA was used to infer a phylogeny using RAxML (-p12345 -# 20 -m PROTGAMMAJTTF) with 100 replications of bootstrapping.

The genomic sequences of 48 additional accessions of *S. pimpinellifolium* were downloaded from NCBI (Supplementary Table 8)[20,40–42]. They were aligned to the contigs of *S. pimpinellifolium* LA1269 using BWA-MEM2 at default settings[96]. The alignments were sorted and indexed using SamTools v1.8[97]. The genes with 45% or more of their exons covered by minimum two reads were considered present by SGSGeneLoss v0.1[43].

### Selection pressure analysis

The pairwise Ks values were calculated for each clade in each species only using intact NLRs. All the pairwise alignments were generated with MAFFT (--maxiterate 1000 --globalpair), and codeML of PAML v4.8[45] was used to estimate the pairwise Ks values using the codon frequency of F3X4. The resulting Ks values were used as distances to hierarchically cluster them with a single linkage. The heights of the dendrograms were used as the representative Ks values.

The global Ka and Ks values were estimated for each OG for the N-terminus, NB-ARC domain and the C-terminus. To define the NB-ARC domain, we collected the NB-ARC domains of all complete NLRs of tomatoes, aligned them using MAFFT (--maxiterate 1000 –globalpair) and trimmed out gappy columns (-gt 0.2). The remaining sequences were used to make a HMM. From the start to the end of the envelope was considered as NB-ARC domain. The global Ka and Ks value were calculated by codeML with the model F3X4.

To detect pervasive and episodic selection pressures, we used FUBAR[98] and MEME[99] in Hyphy v2.3.14.20190429beta(MP)[46]. The default setting was used for FUBAR and MEME, Shannon’s entropy was also calculated using the MSAs. All columns with 40% or more sites were considered for the calculation, and the average Shannon’s entropy value was drawn.

## Acknowledgements

We want to appreciate Freddy Monteiro for helpful discussions and guidance and Ksenia Krasileva for sincere comments and advice. This work was supported by a grant from the Gordon and Betty Moore Foundation to the 2 Blades Foundation (GBMF4725).

---

**Supplementary Figure 1.**
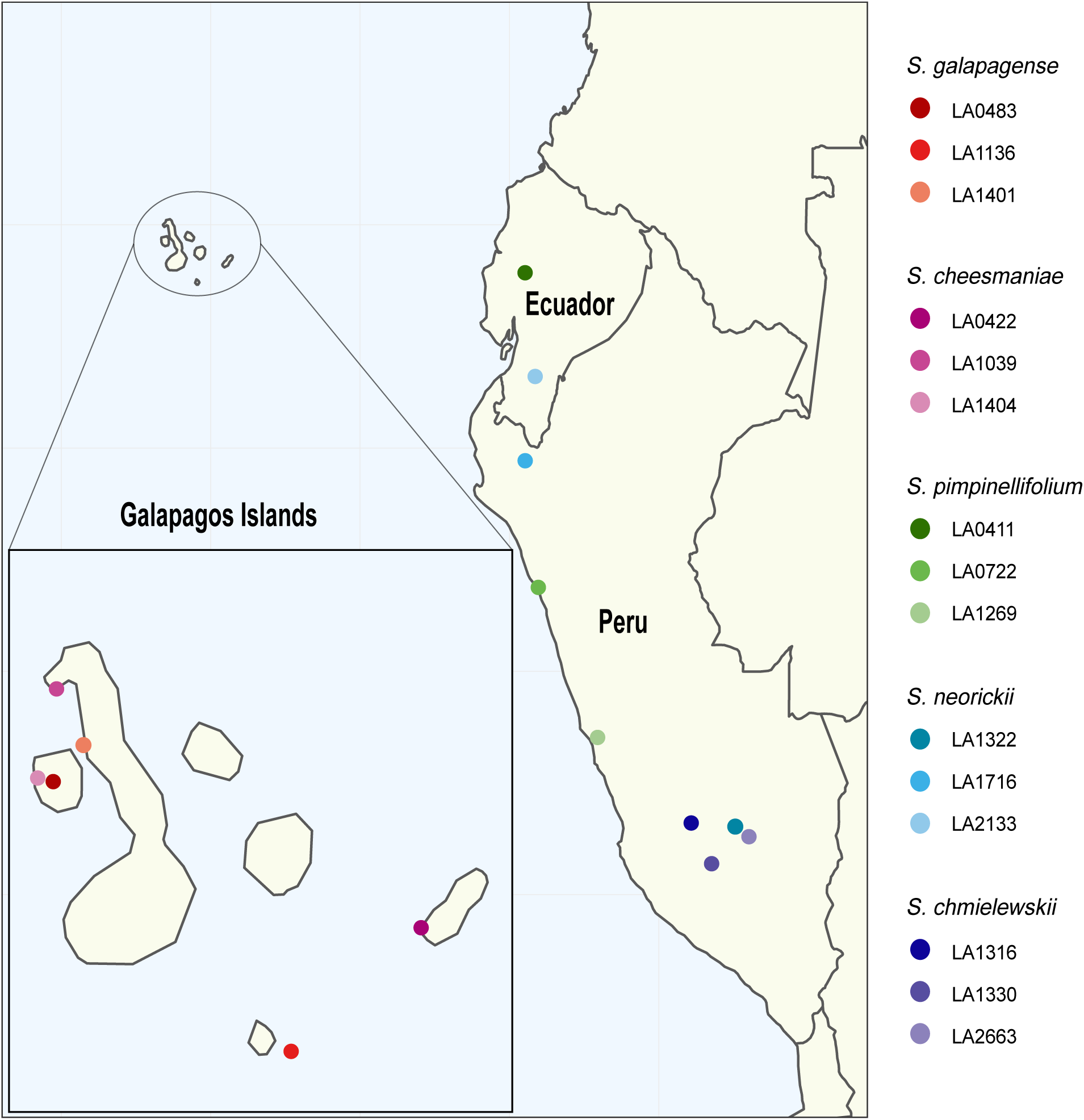
The collection sites of the seeds of the tomatoes used in the study. More information about the accessions can be found in this link: https://tgrc.ucdavis.edu/Data/Acc

**Supplementary Figure 2.**
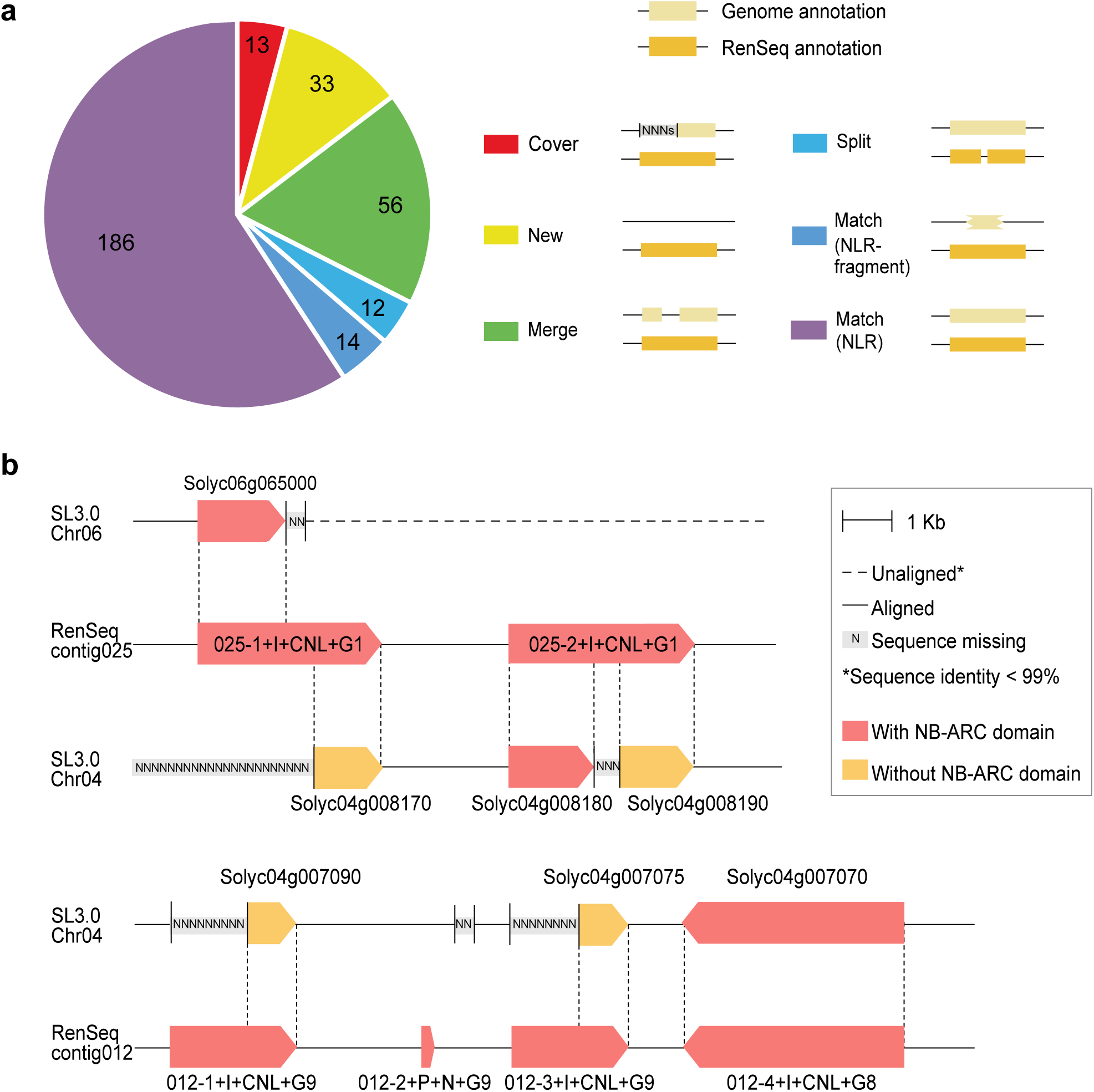
The comparison between ITAG 3.2 and our RenSeq result. The contigs and the genomic sequences of the annotated NLRs from the RenSeq pipeline were mapped to the reference genome (SL 3.0) with a sequence identity of 99% or more as a cut-off. **a.** The 314 annotated NLRs from our RenSeq pipeline were categorized into 6 classes given in the legend in comparison to the annotations of ITAG 3.2. **b.** The examples of the Cover class (red in A) where our RenSeq annotation improved the existing gene models were given.

**Supplementary Figure 3.**
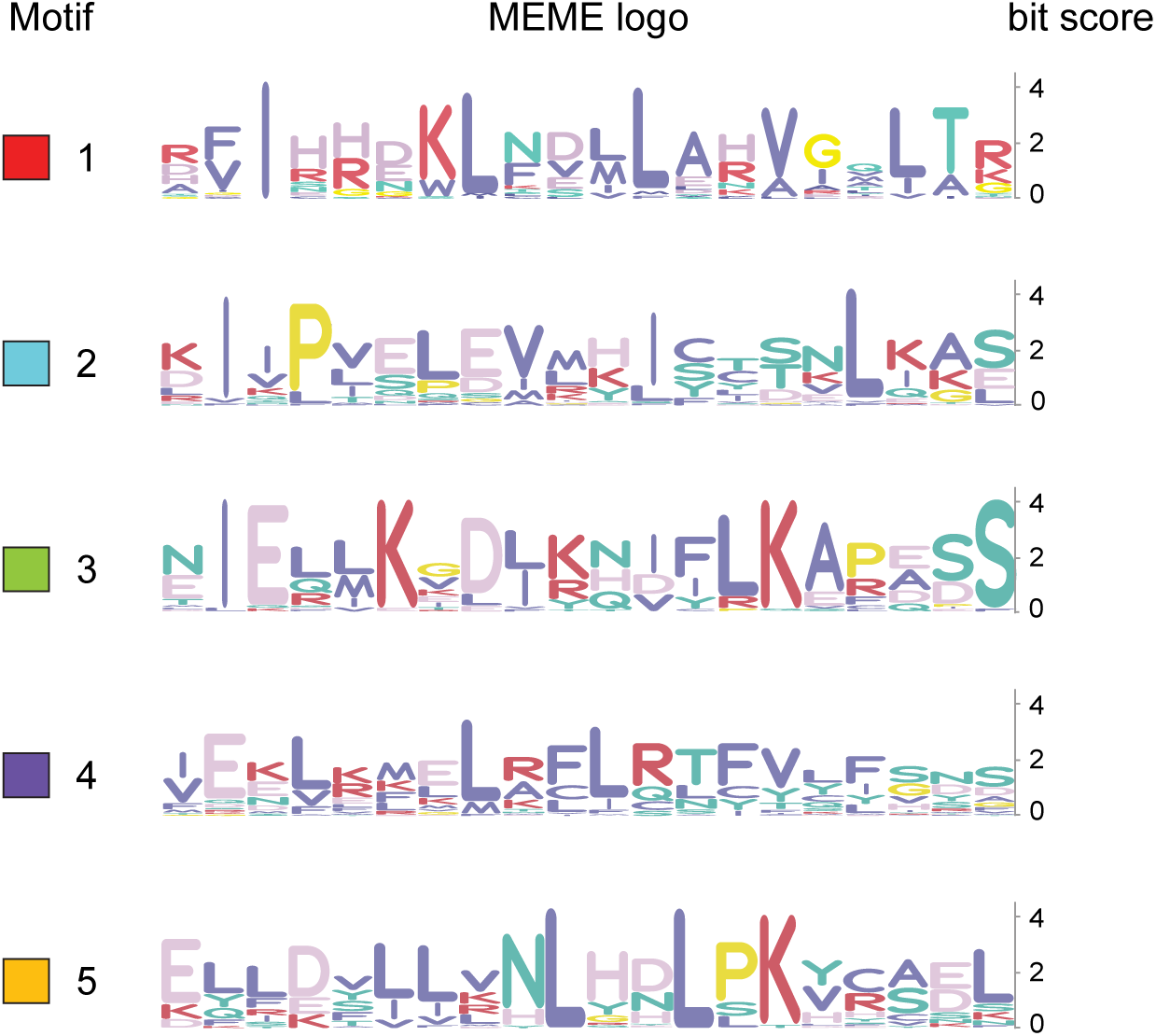
Conserved motifs in the extended N-terminus The five most conserved motifs were identified by MEME in the extended N-terminus sequences of G1, G3 and G6. The annotation of the motifs (the color of the boxes and the numbers) were consistently given in all figures.

**Supplementary Figure 4.**
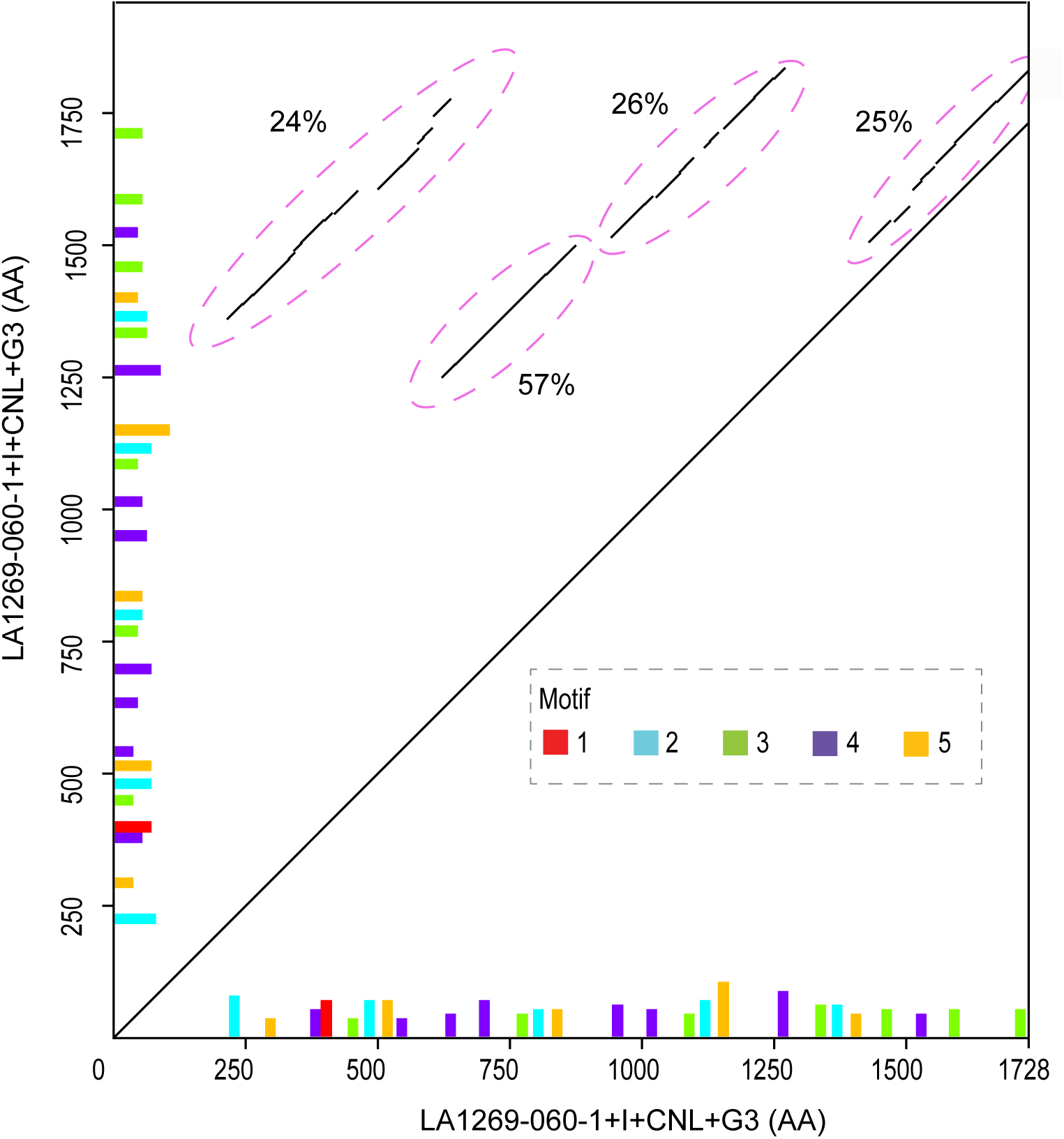
Homology bewteen the motifs in the longest gene model The extended N-terminus sequences of LA1269-060-1+I+CNL+G3 were searched against its entire amino acid sequences. The location of the motifs and the sequence identity of the homologous regions were annotated in the dot plot.

**Supplementary figure 5.**
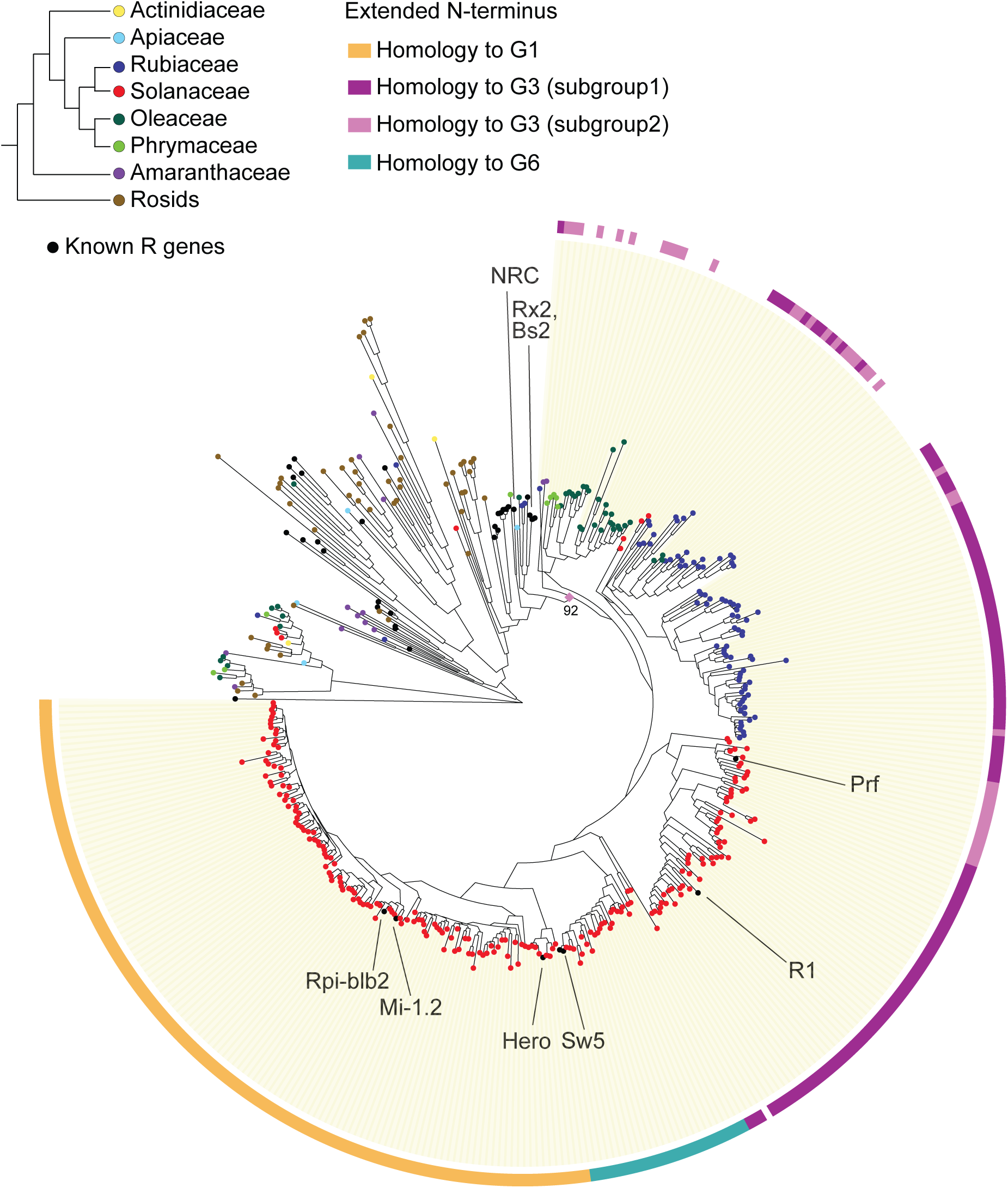
Phylogeny of NB-ARC domains of NLRs with the extended N-terminus NB-ARC domains of NLRs with the extended N-terminus (300 amino acids prior) were used to infer the phylogenetic tree using FastTree with 500 replications of bootsrapping. If the extended N-terminal sequences shared homology with the N-terminal domains of G1, G3 and G6 (E-value < 1E-10), the information was indicated.

**Supplementary Figure 6.**
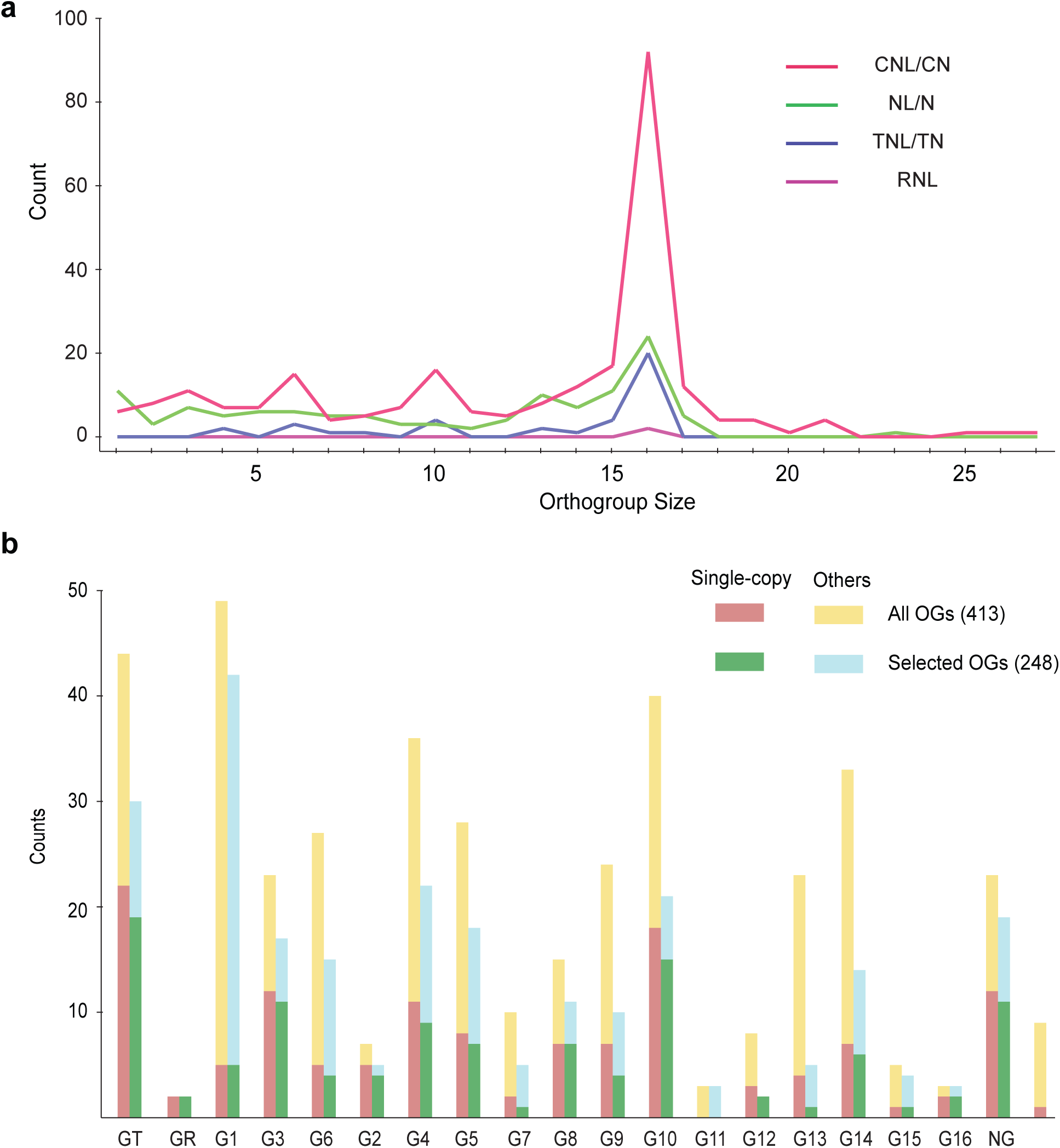
Statistics of curated OGs a. The distribution of the 413 curated OGs based on the size and representative class. b. The number of single-copy OGs and non single-copy OGs for each clade for the 413 curated OGs and 248 selected OGs in which 80% or members have predicted intact NB-ARC domains.

**Supplementary Figure 7.**
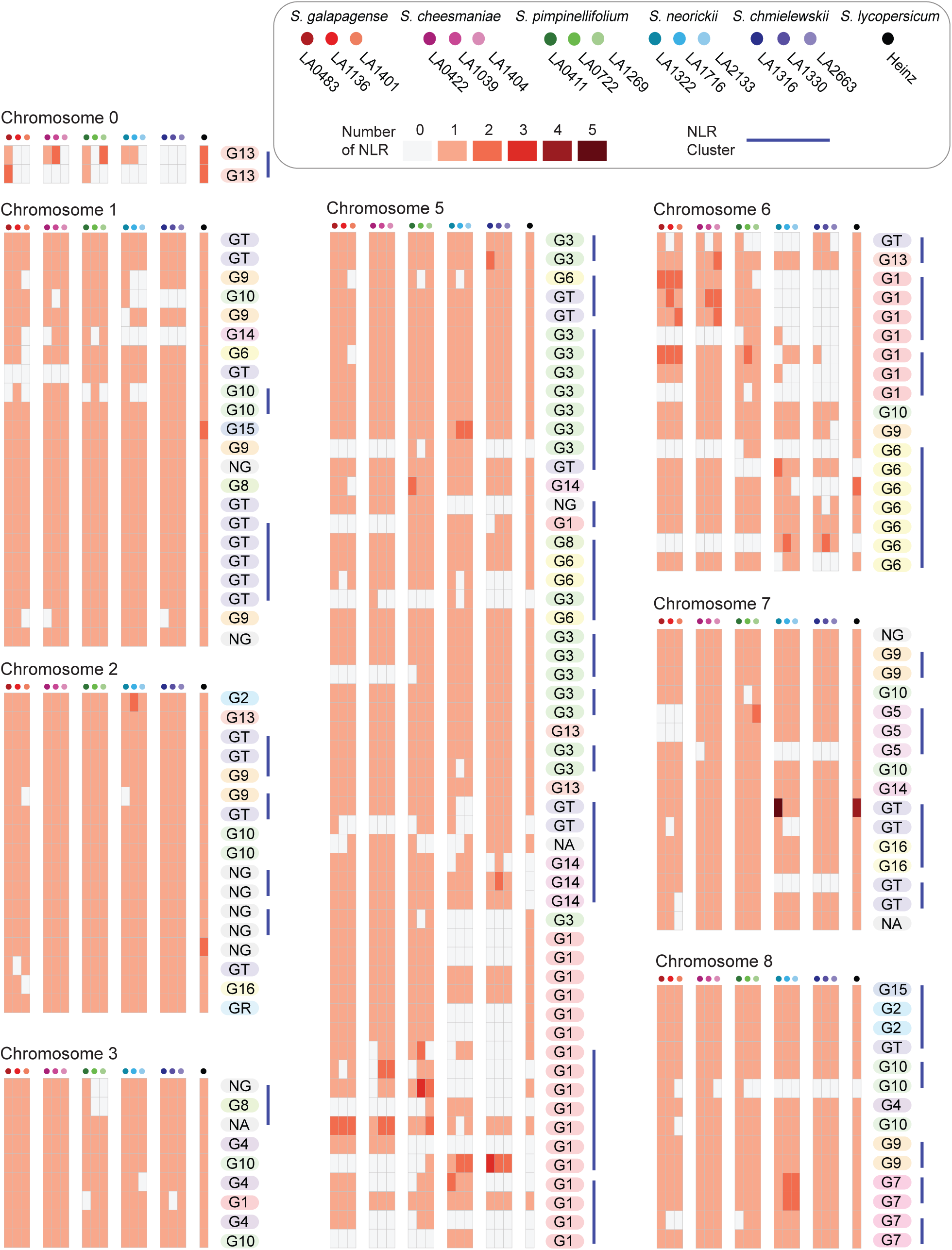
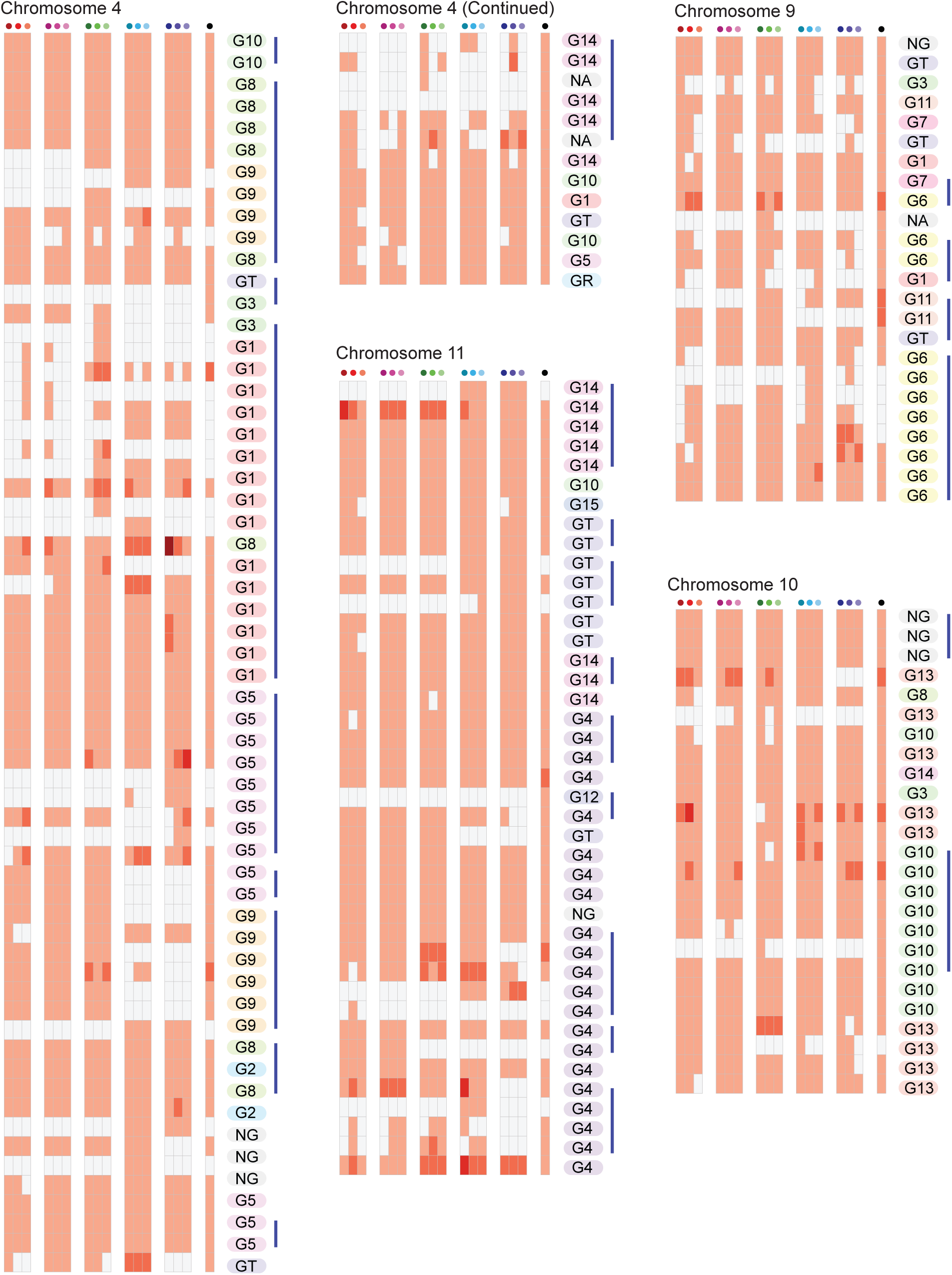
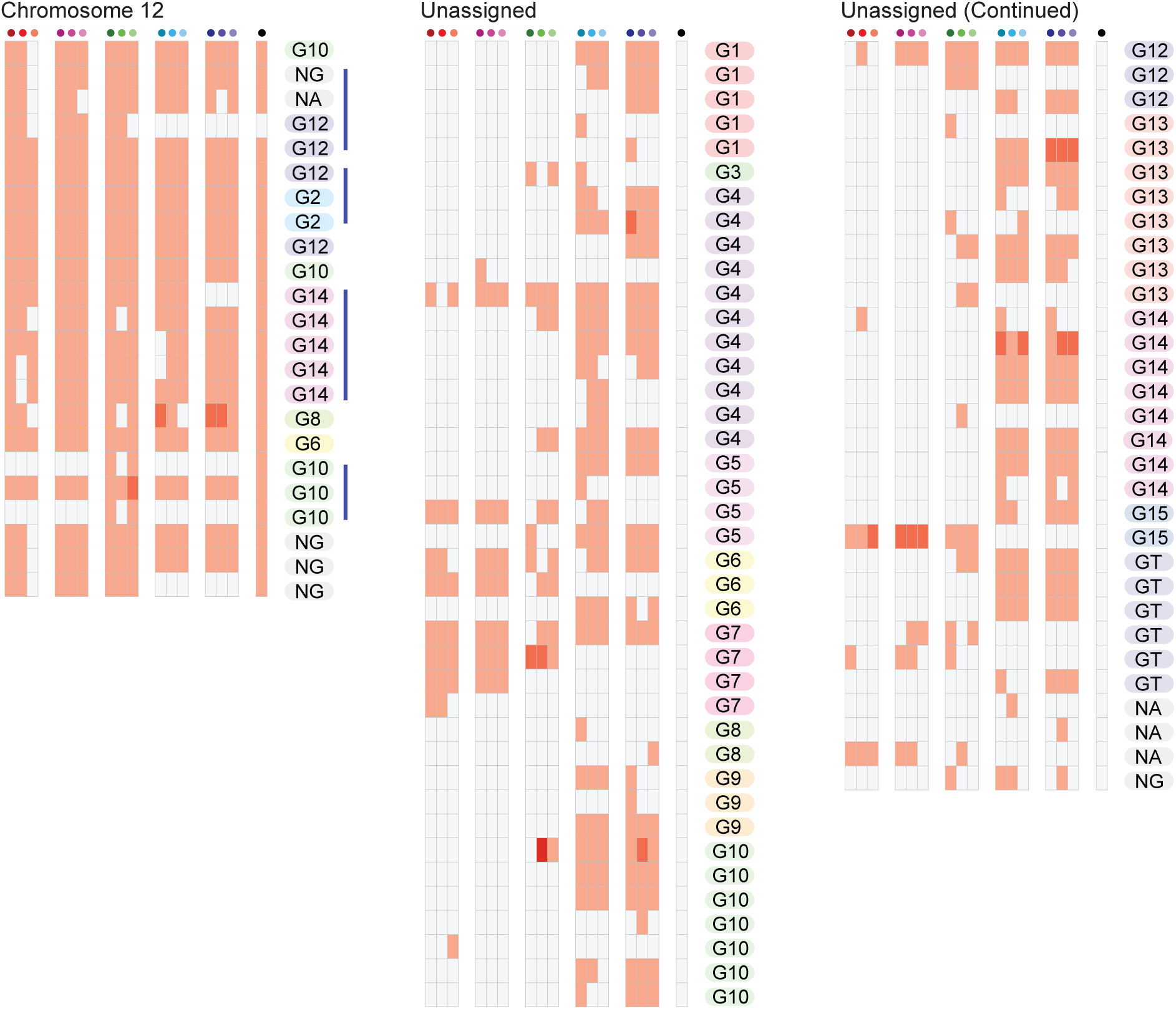
OG-anchoring statistics OGs were anchored to the reference genome. 342 OGs were successfully anchored using NLRs from Heinz and/or the information of NLR gene clusters. For each chromosome, the number of NLRs was indicated with colors for each species. OGs with 200kb or less intergenic regions between were grouped with a blue line, denoting NLR gene clusters. Unanchored OGs were grouped based on clades regardless of their putative loci.

**Supplementary Figure 8.**
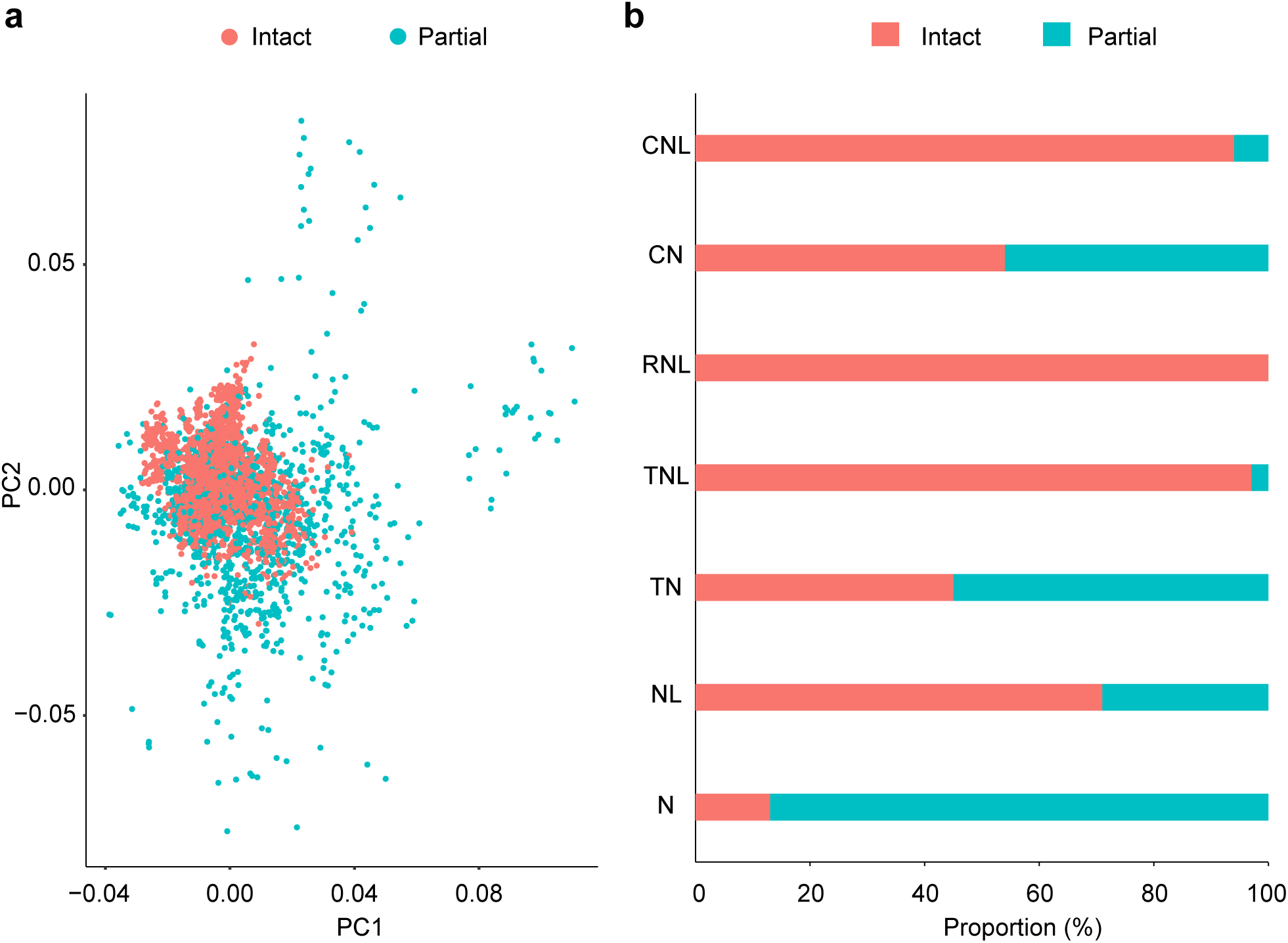
The association of partial NB-ARC domains with the unusual codon usage and incomplete MDAs. NLRs predicted to have intact NB-ARC domains and partial NB-ARC domains were compared. a. Their codon usages were analyzed with PCA and visualized in the two dimensions. b. Their proportions were calculated for each MDA.

**Supplementary Figure 9.**
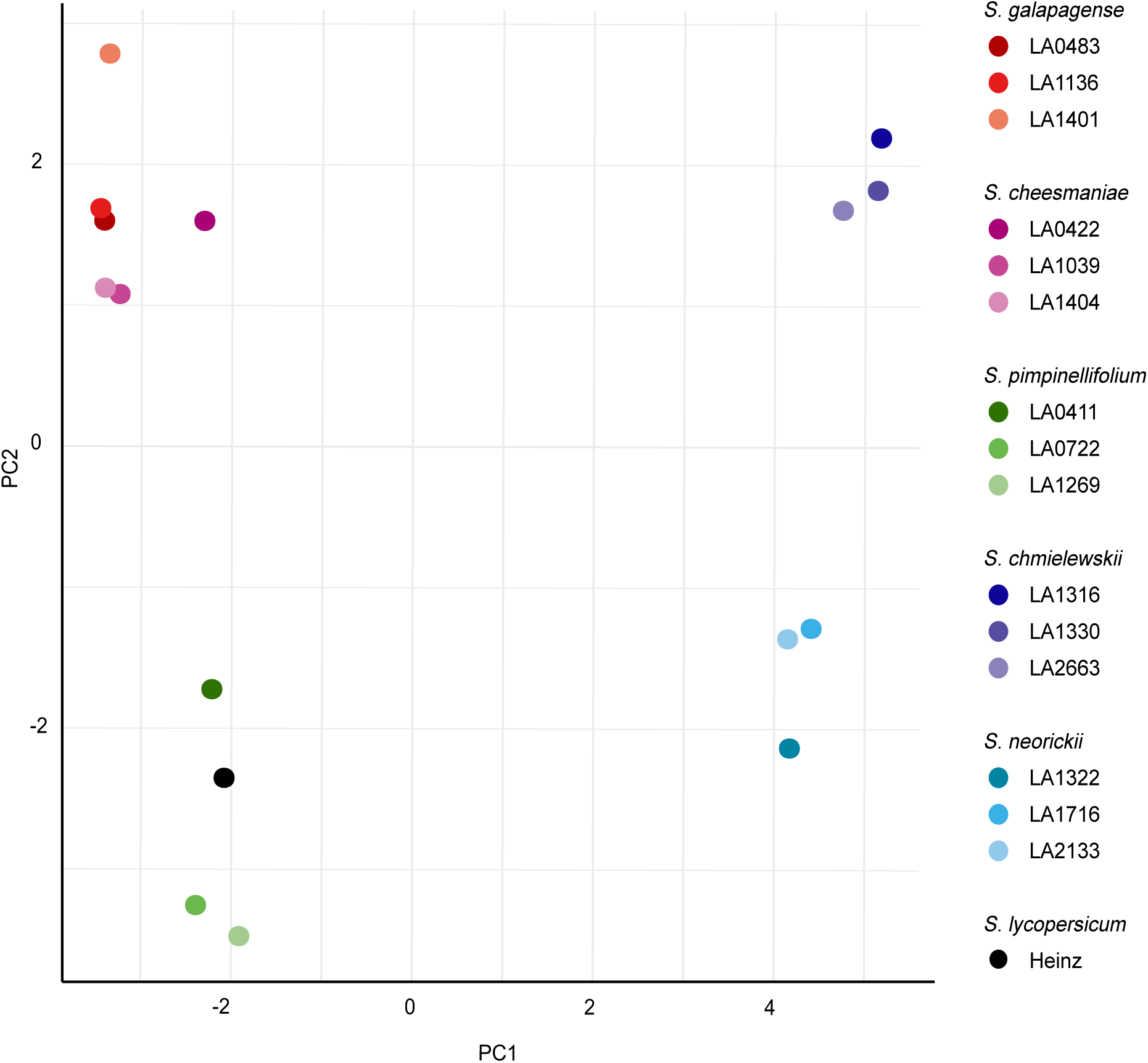
The PCA plot of non-single copy OGs Among the OGs where 80% or more members were classified to contain intact NB-ARC domains, non-single copy OGs were analyzed using PCA. The proximity between points indicates their similarity in the pattern in the OGs.

**Supplementary Figure 10.**
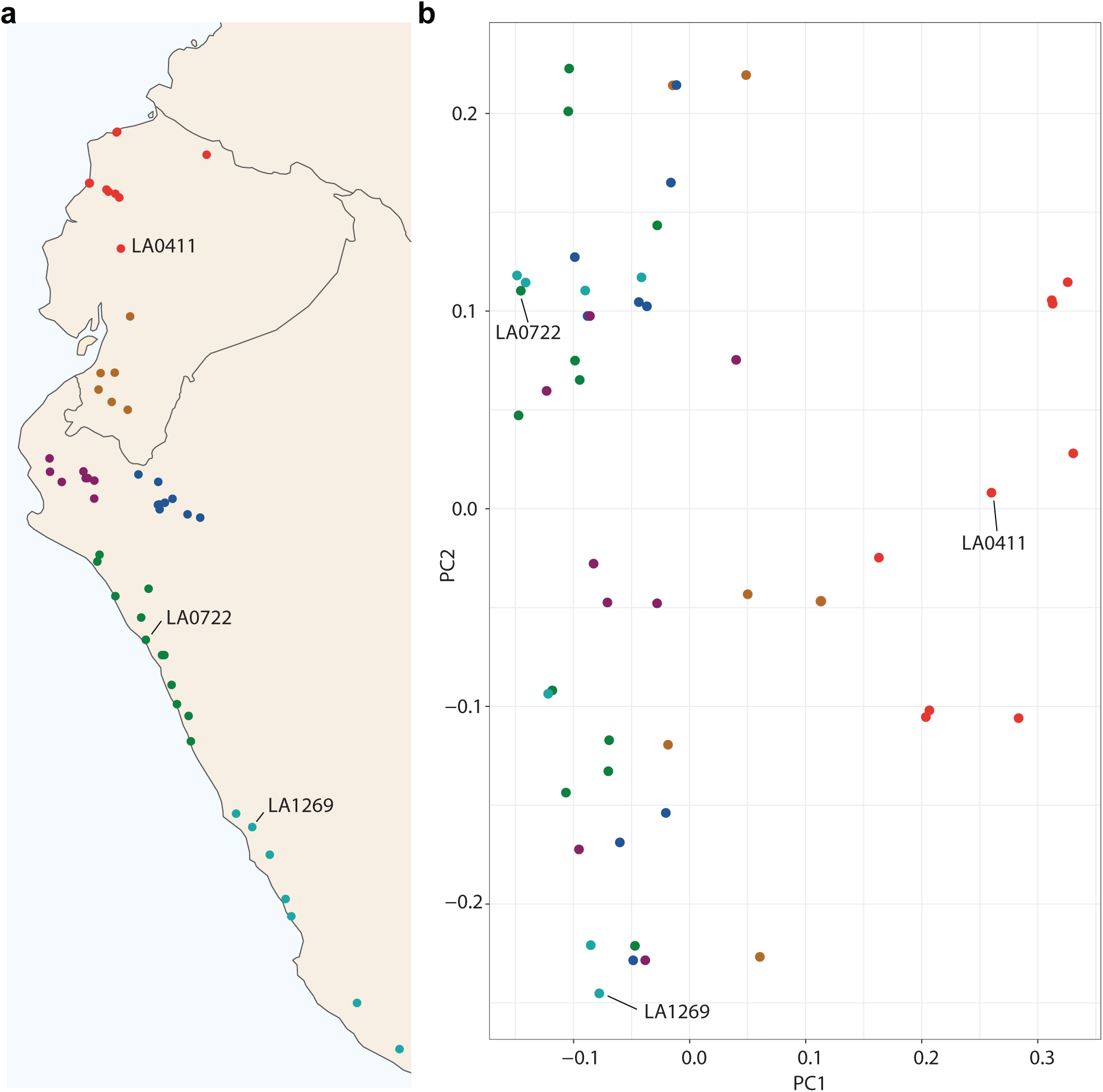
The variations between accessions in *S. pimpinellifolium* The genomic sequences of 48 additional accessions of *S. pimpinelliforlium* were mapped to the contigs of *S. pimpinellifolium* LA1269. The PAV was detected for the OGs in which at least a single gene was for present for LA1269 and 80% or more members were predicted to contain intact NB-ARC domains. a. The collection site of the additional 48 accessions and three accessions used in this study. b. The PCA plot generated with the result of the PAV prediction.

**Supplementary Figure 11.**
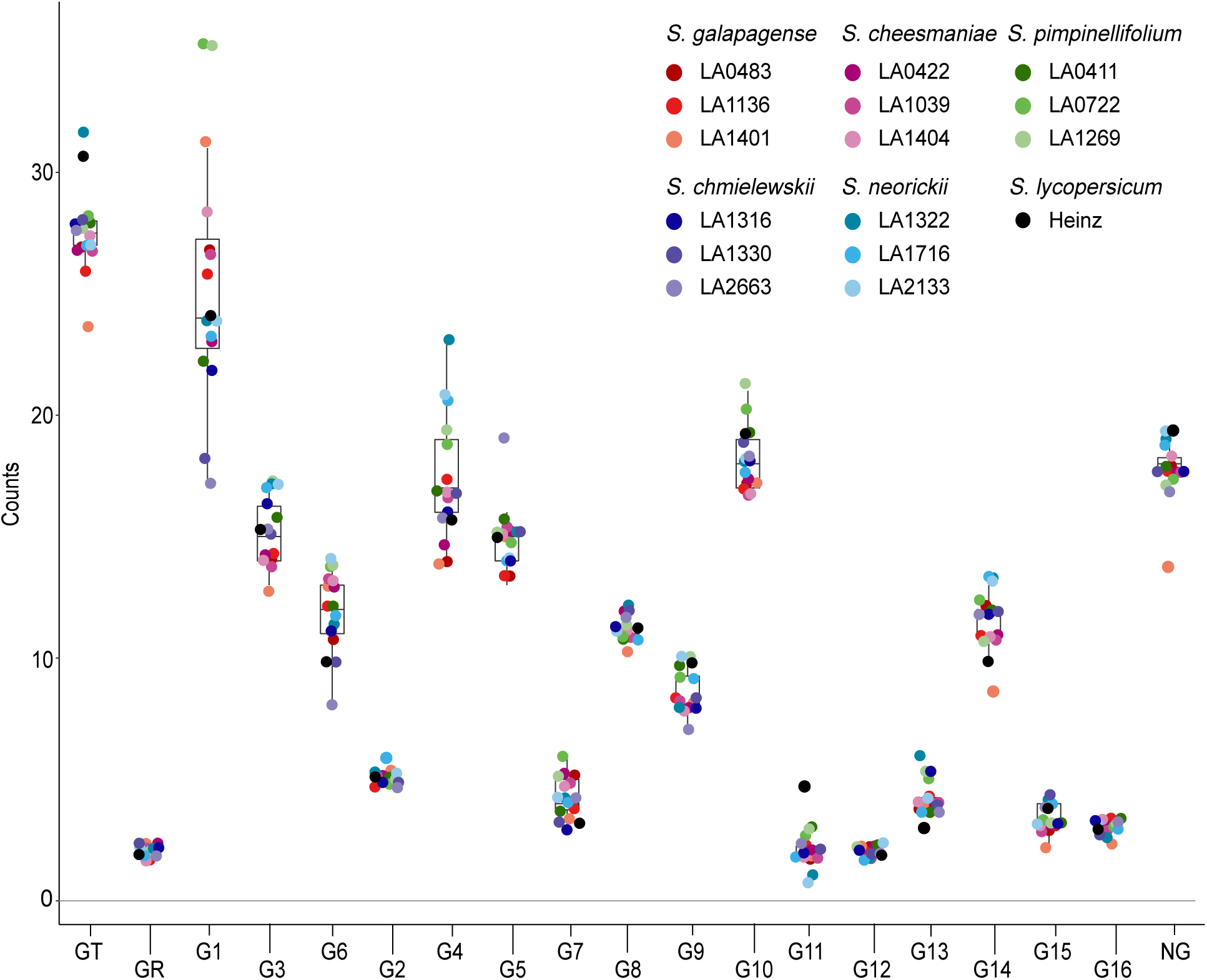
The distribution of NLRs in the 248 selected OGs The number of NLRs in the 248 selected OGs were plotted for each clade for all the accessions in the study.

**Supplementary Figure 12.**
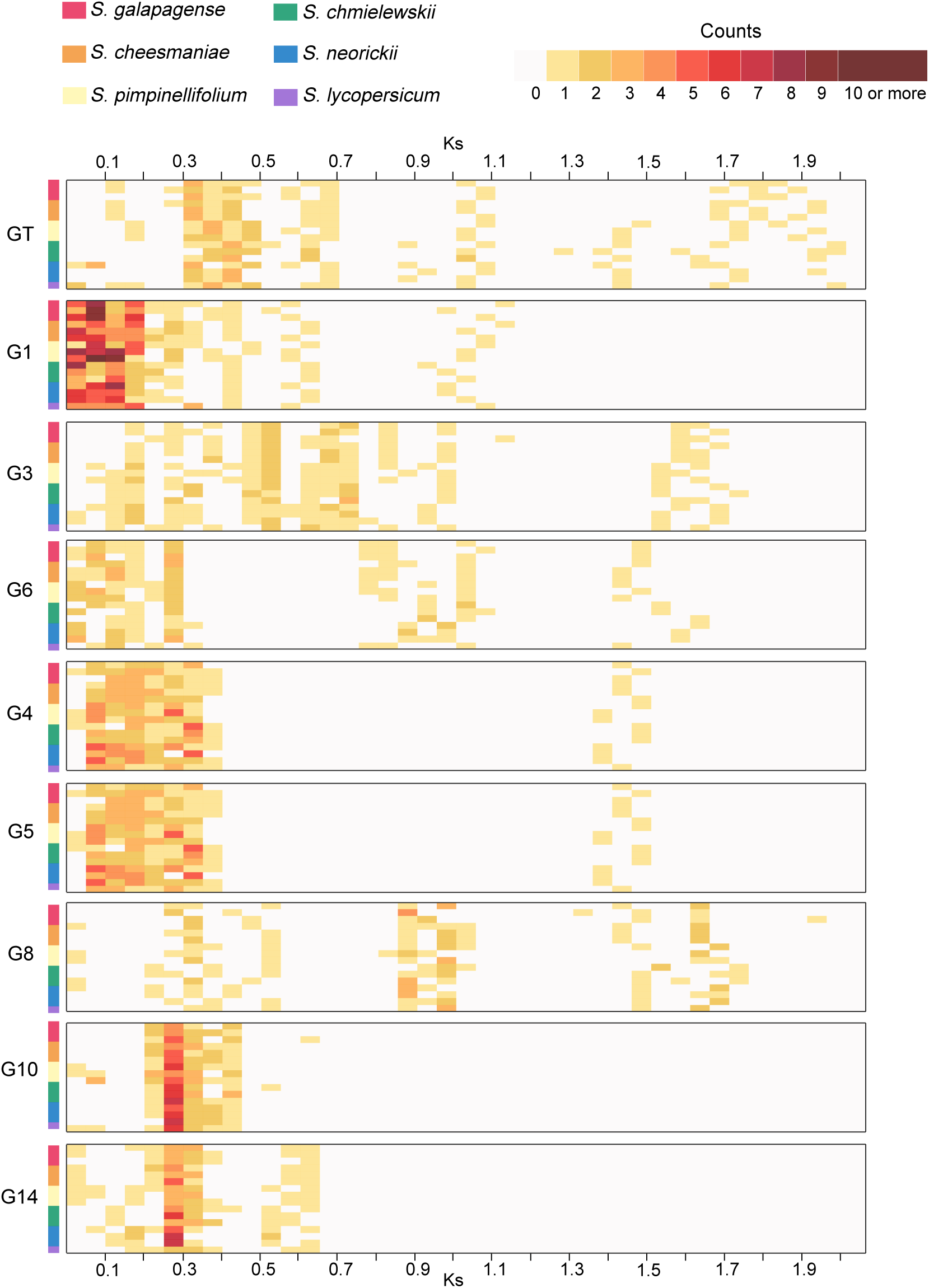
The distribution of the pairwise Ks values The members in the selected OGs where 80% or members were predicted to contain intact NB-ARC domains were used to compare their pairwise Ks values for each calde and for each species. Representative Ks values were calculated using hierarchical clustering, and their distributions were visualized as heatmaps.

**Supplementary Figure 13.**
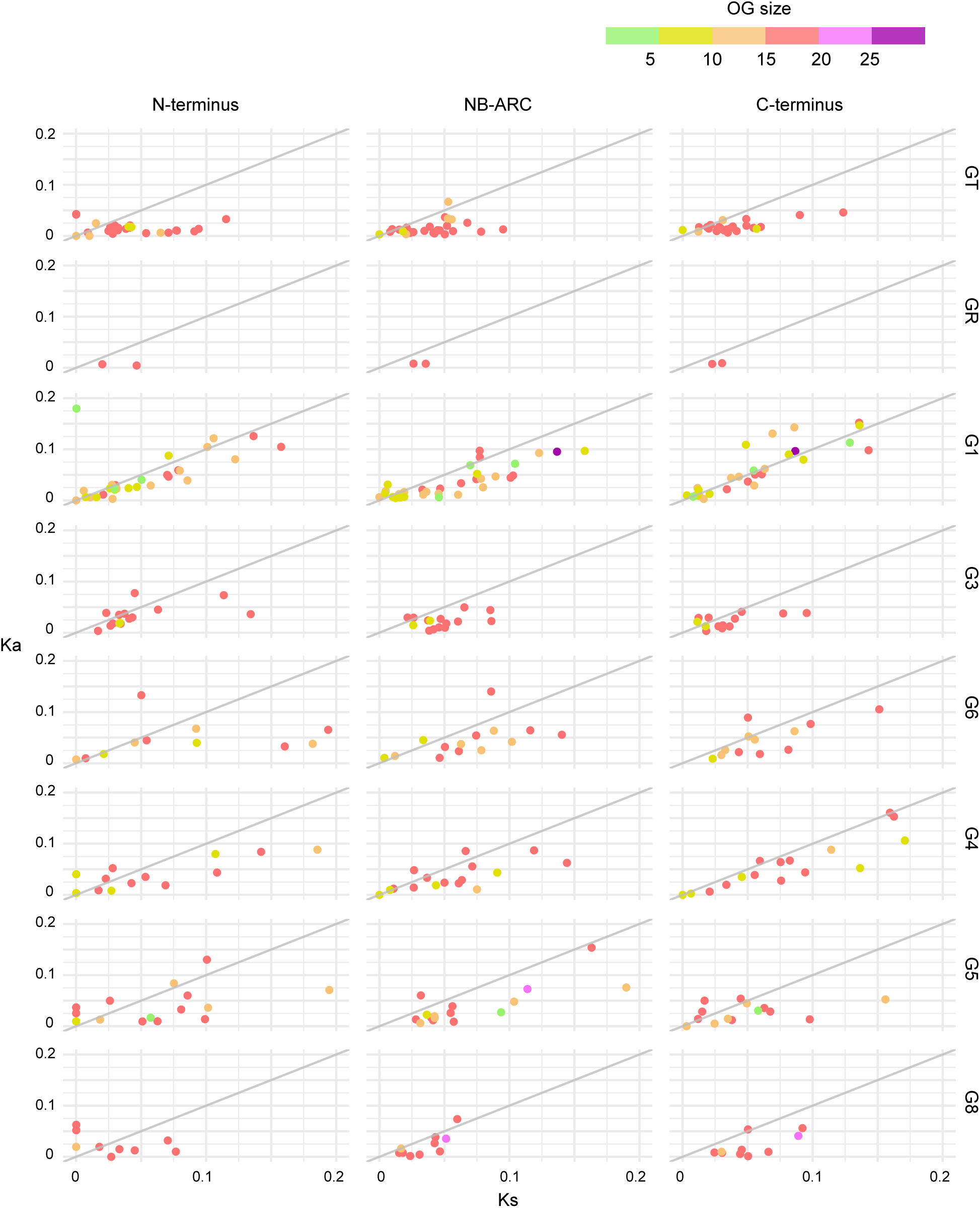
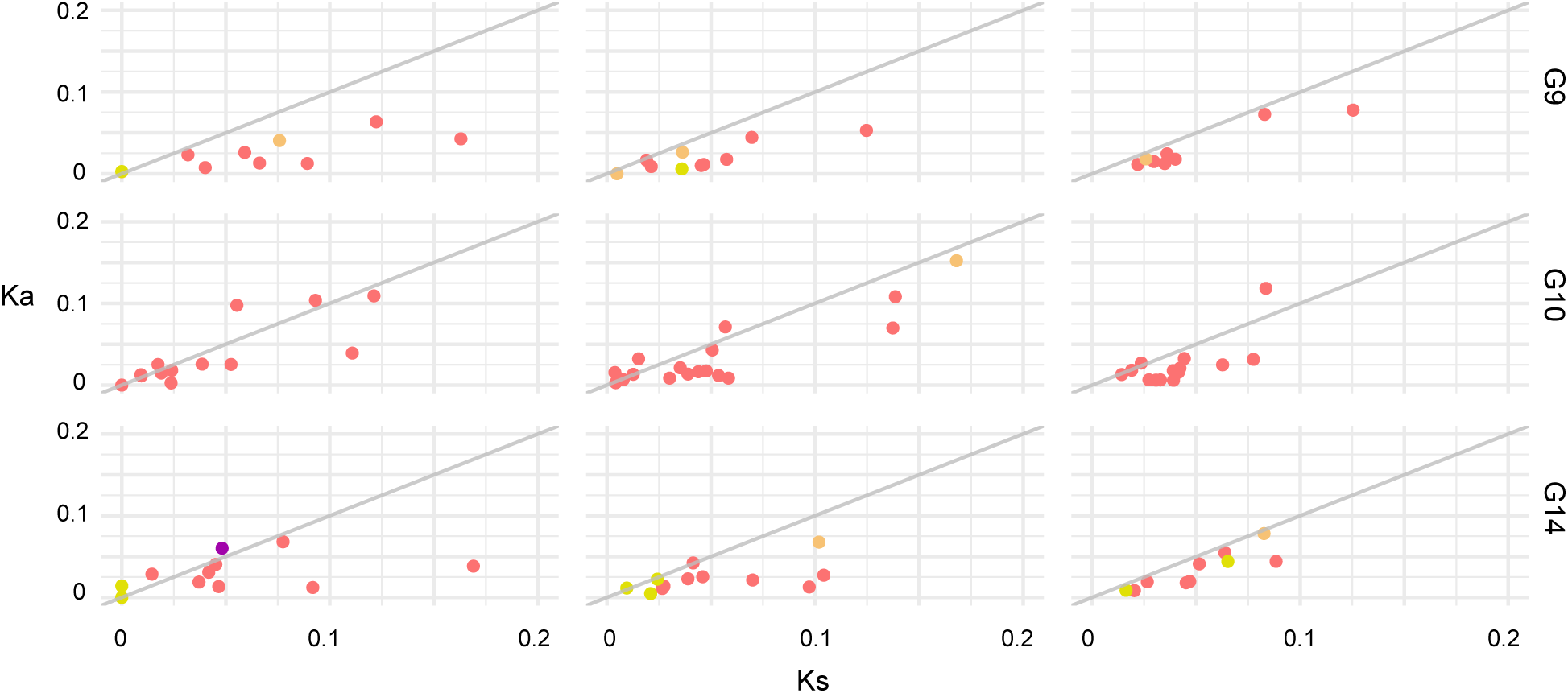
Global Ka and Ks values for OGs The selected OGs in which 80% or members were predicted to contain intact NB-ARC domains were aligned with Prank. For each OG, global Ka and Ks values were calculated using CodeML in PAML. The OGs were classified based on their size and clades, and their Ka and Ks values were plotted.

## Supporting information

Supplementary Table 1. Plant materials.

Supplementary Table 2. Sequencing statistics.

Supplementary Table 3. The number of NLRs based on MDA.

Supplementary Table 4. The comparison of NLRs in *N. benthamiana*.

Supplementary Table 5. The number of NLRs based on phylogenetic relationship.

Supplementary Table 6. Comparative analysis of extended NLRs.

Supplementary Table 7. The list of orthogroups.

Supplementary Table 8. *S. pimpinellifolium* NLR comparison source.

Supplementary Table 9. The number of NLRs based on orthogroups.

## References

1. Dangl JL, Horvath DM, Staskawicz BJ. Pivoting the plant immune system from dissection to deployment. Science (80-). 2013;341:746–51.

2. Jones JD, Dangl JL. The plant immune system. Nature. 2006;444:323–9.

3. Shao Z-Q, Xue J-Y, Wu P, Zhang Y-M, Wu Y, Hang Y-Y, et al. Large-scale analyses of angiosperm nucleotide-binding site-leucine-rich repeat genes reveal three anciently diverged classes with distinct evolutionary patterns. Plant Physiol. 2016;170:1095–2109.

4. Sarris PF, Cevik V, Dagdas G, Jones JDG, Krasileva K V. Comparative analysis of plant immune receptor architectures uncovers host proteins likely targeted by pathogens. BMC Biol. BMC Biology; 2016;14:8.

5. Ellis JG. Integrated decoys and effector traps: how to catch a plant pathogen. BMC Biol. 2016;14:13.

6. Mucyn TS, Clemente A, Andriotis VM, Balmuth AL, Oldroyd GE, Staskawicz BJ, et al. The tomato NBARC-LRR protein Prf interacts with Pto kinase in vivo to regulate specific plant immunity. Plant Cell. 2006;18:2792–806.

7. Lukasik-Shreepaathy E, Slootweg E, Richter H, Goverse A, Cornelissen BJ, Takken FL. Dual regulatory roles of the extended N terminus for activation of the tomato MI-1.2 resistance protein. Mol Plant Microbe Interact. 2012;25:1045–57.

8. Li J, Huang H, Zhu M, Huang S, Zhang W, Dinesh-Kumar SP, et al. A plant immune receptor adopts a two-step recognition mechanism to enhance viral effector perception. Mol Plant. Elsevier Ltd; 2019;12:248–62.

9. Zheng F, Wu H, Zhang R, Li S, He W, Wong FL, et al. Molecular phylogeny and dynamic evolution of disease resistance genes in the legume family. BMC Genomics. 2016;17:402.

10. Mondragón-Palomino M, Stam R, John-Arputharaj A, Dresselhaus T. Diversification of defensins and NLRs in Arabidopsis species by different evolutionary mechanisms. BMC Evol Biol. 2017;17:255.

11. Leister D. Tandem and segmental gene duplication and recombination in the evolution of plant disease resistance gene. Trends Genet. 2004;20:116–22.

12. Zhong Y, Yin H, Sargent DJ, Malnoy M, Cheng ZM. Species-specific duplications driving the recent expansion of NBS-LRR genes in five Rosaceae species. BMC Genomics. 2015;16:77.

13. Zhang R, Murat F, Pont C, Langin T, Salse J. Paleo-evolutionary plasticity of plant disease resistance genes. BMC Genomics. 2014;15:187.

14. Seo E, Kim S, Yeom S-I, Choi D. Genome-wide comparative analyses reveal the dynamic evolution of nucleotide-binding leucine-rich repeat gene family among Solanaceae plants. Front Plant Sci. 2016;7:1–13.

15. Kim S, Park J, Yeom SI, Kim YM, Seo E, Kim KT, et al. New reference genome sequences of hot pepper reveal the massive evolution of plant disease-resistance genes by retroduplication. Genome Biol. Genome Biology; 2017;18:1–11.

16. Wu C-H, Abd-El-Haliem A, Bozkurt TO, Belhaj K, Terauchi R, Vossen JH, et al. NLR network mediates immunity to diverse plant pathogens. Proc Natl Acad Sci. 2017;114:8113–8.

17. Wu C-H, Derevnina L, Kamoun S. Receptor networks underpin plant immunity. Science (80-). 2018;360:1300–1.

18. Lukyanenko AN. Disease Resistance in Tomato. Genet Improv Tomato. 1991. p. 99–119.

19. Rick CM, Chetelat RT. Utilization of related wild species for tomato improvement. Acta Hortic. 1995;412.

20. Gao L, Gonda I, Sun H, Ma Q, Bao K, Tieman DM, et al. The tomato pan-genome uncovers new genes and a rare allele regulating fruit flavor. Nat Genet. 2019;51:1044–51.

21. Walter JM. Hereditary resistance to disease in tomato. Annu Rev Phytopathol. 1967;5:131–60.

22. Peralta IE, Knapp S, Spooner DM. New species of wild tomatoes (Solanum section Lycopersicon: Solanaceae) from Northern Peru. Syst Bot. 2005;30:424–34.

23. Bergougnoux V. The history of tomato: From domestication to biopharming. Biotechnol Adv. 2014;32:170–89.

24. Chaudhary R, Atamian H. Resistance-Gene-Mediated Defense Responses against Biotic Stresses in the Crop Model Plant Tomato. J Plant Pathol Microbiol. 2017;8:4.

25. Fukuoka S, Saka N, Mizukami Y, Koga H, Yamanouchi U, Yoshioka Y, et al. Gene pyramiding enhances durable blast disease resistance in rice. Sci Rep. 2015;5:1–7.

26. Ghislain M, Byarugaba AA, Magembe E, Njoroge A, Rivera C, Román ML, et al. Stacking three late blight resistance genes from wild species directly into African highland potato varieties confers complete field resistance to local blight races. Plant Biotechnol J. 2019;17:1119–29.

27. Bauchet G, Causse M. Genetic diversity in tomato (Solanum lycopersicum) and its wild relatives. Genet Divers Plants. 2012. p. 133–62.

28. Stam R, Silva-Arias GA, Tellier A. Subsets of NLR genes show differential signatures of adaptation during colonisation of new habitats. New Phytol. 2019;224:367–79.

29. Jupe F, Witek K, Verweij W, Sliwka J, Pritchard L, Etherington GJ, et al. Resistance gene enrichment sequencing (RenSeq) enables reannotation of the NB-LRR gene family from sequenced plant genomes and rapid mapping of resistance loci in segregating populations. Plant J. 2013;76:530–44.

30. Andolfo G, Jupe F, Witek K, Etherington GJ, Ercolano MR, Jones JD. Defining the full tomato NB-LRR resistance gene repertoire using genomic and cDNA RenSeq. BMC Plant Biol. 2014;14:120.

31. Witek K, Jupe F, Witek AI, Baker D, Clark MD, Jones JDG. Accelerated cloning of a potato late blight-resistance gene using RenSeq and SMRT sequencing. Nat Biotechnol. 2016;34:656–60.

32. Van de Weyer A-L, Monteiro F, Furzer OJ, Nishimura MT, Cevik V, Witek K, et al. A species-wide inventory of NLR genes and alleles in Arabidopsis thaliana. Cell. Elsevier Inc.; 2019;178:1260–1272.e14.

33. Bombarely A, Rosli HG, Vrebalov J, Moffett P, Mueller LA, Martin GB. A daft genome sequence of Nicotiana benthamiana to enhance molecular plant-microbe biology research. Mol Plant-Microbe Interact. 2012;25:1523–30.

34. Naim F, Nakasugi K, Crowhurst RN, Hilario E, Zwart AB, Hellens RP, et al. Advanced engineering of lipid metabolism in Nicotiana benthamiana using a draft genome and the V2 viral silencing-suppressor protein. PLoS One. 2012;7:e52717.

35. Steuernagel B, Jupe F, Witek K, Jones JDG, Wulff BBH. NLR-parser: Rapid annotation of plant NLR complements. Bioinformatics. 2015;31:1665–7.

36. Eddy SR. Profile hidden Markov models. Bioinformatics. 1998;14:755–63.

37. Bailey TL, Boden M, Buske FA, Frith M, Grant CE, Clementi L, et al. MEME SUITE: tools for motif discovery and searching. Nucleic Acids Res. 2009;37:W202–8.

38. Emms DM, Kelly S. OrthoFinder: solving fundamental biases in whole genome comparisons dramatically improves orthogroup inference accuracy. Genome Biol. 2015;16:157.

39. Krasileva K V., Buffalo V, Bailey P, Pearce S, Ayling S, Tabbita F, et al. Separating homeologs by phasing in the tetraploid wheat transcriptome. Genome Biol. 2013;14:R66.

40. Aflitos S, Schijlen E, De Jong H, De Ridder D, Smit S, Finkers R, et al. Exploring genetic variation in the tomato (Solanum section Lycopersicon) clade by whole-genome sequencing. Plant J. 2014;80:136– 48.

41. Lin T, Zhu G, Zhang J, Xu X, Yu Q, Zheng Z, et al. Genomic analyses provide insights into the history of tomato breeding. Nat Genet. 2014;46:1220–6.

42. The Tomato Genome Consortium. The tomato genome sequence provides insights into fleshy fruit evolution. Nature. 2012;485:635–41.

43. Golicz AA, Martinez PA, Zander M, Patel DA, Van De Wouw AP, Visendi P, et al. Gene loss in the fungal canola pathogen Leptosphaeria maculans. Funct Integr Genomics. 2015;15:189–96.

44. Kim S, Park M, Yeom S-I, Kim Y-M, Lee JM, Lee H-A, et al. Genome sequence of the hot pepper provides insights into the evolution of pungency in Capsicum species. Nat Genet. 2014;46:270–8.

45. Yang Z. PAML 4: Phylogenetic analysis by maximum likelihood. Mol Biol Evol. 2007;24:1586–91.

46. Kosakovsky Pond SL, Frost SDW, Muse S V. HyPhy: Hypothesis testing using phylogenies. Bioinformatics. 2005;21:676–9.

47. Bai Y, Lindhout P. Domestication and breeding of tomatoes: What have we gained and what can we gain in the future? Ann Bot. 2007;100:1085–94.

48. Björklund ÅK, Ekman D, Elofsson A. Expansion of protein domain repeats. PLoS Comput Biol. 2006;2:e114.

49. Pedley KF, Martin GB. Molecular basis of Pto-mediated resistance to bacterial speck disease in tomato. Annu Rev Phytopathol. 2003;41:215–43.

50. Mucyn TS, Wu AJ, Balmuth AL, Arasteh JM, Rathjen JP. Regulation of tomato Prf by Pto-like protein kinases. Mol Plant-Microbe Interact. 2009;22:391–401.

51. Gutierrez JR, Balmuth AL, Ntoukakis V, Mucyn TS, Gimenez-Ibanez S, Jones AME, et al. Prf immune complexes of tomato are oligomeric and contain multiple Pto-like kinases that diversify effector recognition. Plant J. 2010;61:507–18.

52. Ntoukakis V, Balmuth AL, Mucyn TS, Gutierrez JR, Jones AME, Rathjen JP. The tomato Prf complex is a molecular trap for bacterial effectors based on Pto tansphosphorylation. PLoS Pathog. 2013;9:e1003123.

53. Baggs E, Dagdas G, Krasileva K V. NLR diversity, helpers and integrated domains: making sense of the NLR IDentity. Curr Opin Plant Biol. 2017;38:59–67.

54. Vos P, Simons G, Jesse T, Wijbrandi J, Heinen L, Hogers R, et al. The tomato Mi-1 gene confers resistance to both root-knot nematodes and potato aphids. Nat Biotechnol. 1998;16:1365–9.

55. Nombela G, Williamson VM, Muñiz M. The root-knot nematode resistance gene Mi-1.2 of tomato is responsible for resistance against the whitefly Bemisia tabaci. Mol Plant-Microbe Interact. 2003;16:645– 9.

56. Casteel CL, Walling LL, Paine TD. Behavior and biology of the tomato psyllid, Bactericerca cockerelli, in response to the Mi-1.2 gene. Entomol Exp Appl. 2006;121:67–72.

57. Ellis PR, Maxon Smith JW. Inheritance of resistance to potato cyst-eelworm (Heterodera rostochiensisWoll.) in the genus Lycopersicon. Euphytica. 1971;20:93–101.

58. Ernst K, Kumar A, Kriseleit D, Kloos DU, Phillips MS, Ganal MW. The broad-spectrum potato cyst nematode resistance gene (Hero) from tomato is the only member of a large gene family of NBS-LRR genes with an unusual amino acid repeat in the LRR region. Plant J. 2002;31:127–36.

59. Schultink A, Qi T, Lee A, Steinbrenner AD, Staskawicz B. Roq1 mediates recognition of the Xanthomonas and Pseudomonas effector proteins XopQ and HopQ1. Plant J. 2017;92:787–95.

60. Qi T, Seong K, Thomazella DPT, Kim JR, Pham J, Seo E, et al. NRG1 functions downstream of EDS1 to regulate TIR-NLR-mediated plant immunity in Nicotiana benthamiana. Proc Natl Acad Sci U S A. 2018;115:E10979–87.

61. Brommonschenkel SH, Frary A, Frary A, Tanksley SD. The broad-spectrum tospovirus resistance gene Sw-5 of tomato is a homolog of the root-knot nematode resistance gene Mi. Mol Plant-Microbe Interact. 2000;13:1130–8.

62. Peiró A, Cañizares MC, Rubio L, López C, Moriones E, Aramburu J, et al. The movement protein (NSm) of Tomato spotted wilt virus is the avirulence determinant in the tomato Sw-5 gene-based resistance. Mol Plant Pathol. 2014;15:802–13.

63. Chen X, Zhu M, Jiang L, Zhao W, Li J, Wu J, et al. A multilayered regulatory mechanism for the autoinhibition and activation of a plant CC-NB-LRR resistance protein with an extra N-terminal domain. New Phytol. 2016;212:161–75.

64. Ye J, Liu G, Chen W, Zhang F, Li H, Ye Z, et al. Knockdown of SlNL33 accumulates ascorbate, enhances disease and oxidative stress tolerance in tomato (Solanum lycopersicum). Plant Growth Regul. 2019;89:49–58.

65. Long M, Betrán E, Thornton K, Wang W. The origin of new genes: Glimpses from the young and old. Nat Rev Genet. 2003;4:865–75.

66. Zhong Y, Cheng ZM. A unique RPW8-encoding class of genes that originated in early land plants and evolved through domain fission, fusion, and duplication. Sci Rep. 2016;6:32923.

67. Wang J, Hu M, Wang J, Qi J, Han Z, Wang G, et al. Reconstitution and structure of a plant NLR resistosome conferring immunity. Science (80-). 2019;364:eaav5870.

68. Camacho C, Coulouris G, Avagyan V, Ma N, Papadopoulos J, Bealer K, et al. BLAST+: architecture and applications. BMC Bioinformatics. 2009;10:421.

69. Martin M. Cutadapt removes adapter sequences from high-throughput sequencing reads. EMBnet.journal. 2011;17:1.

70. Koren S, Walenz BP, Berlin K, Miller JR, Bergman NH, Phillippy AM. Canu: Scalable and accurate long-read assembly via adaptive κ-mer weighting and repeat separation. Genome Res. 2017;27:722–36.

71. Gurevich A, Saveliev V, Vyahhi N, Tesler G. QUAST: Quality assessment tool for genome assemblies. Bioinformatics. 2013;29:1072–5.

72. Cantarel BL, Korf I, Robb SMC, Parra G, Ross E, Moore B, et al. MAKER: An easy-to-use annotation pipeline designed for emerging model organism genomes. Genome Res. 2008;18:188–96.

73. Stanke M, Schöffmann O, Morgenstern B, Waack S. Gene prediction in eukaryotes with a generalized hidden Markov model that uses hints from external sources. BMC Bioinformatics. 2006;7:62.

74. Korf I. Gene finding in novel genomes. BMC Bioinformatics. 2004;5:59.

75. Li W, Godzik A. Cd-hit: A fast program for clustering and comparing large sets of protein or nucleotide sequences. Bioinformatics. 2006;22:1658–9.

76. Lomsadze A, Ter-Hovhannisyan V, Chernoff YO, Borodovsky M. Gene identification in novel eukaryotic genomes by self-training algorithm. Nucleic Acids Res. 2005;33:6494–506.

77. Jones P, Binns D, Chang HY, Fraser M, Li W, McAnulla C, et al. InterProScan 5: Genome-scale protein function classification. Bioinformatics. 2014;30:1236–40.

78. Finn RD, Bateman A, Clements J, Coggill P, Eberhardt RY, Eddy SR, et al. Pfam: The protein families database. Nucleic Acids Res. 2014;42:D222–30.

79. Bolger A, Scossa F, Bolger ME, Lanz C, Maumus F, Tohge T, et al. The genome of the stress-tolerant wild tomato species Solanum pennellii. Nat Genet. Nature Publishing Group; 2014;46:1034–8.

80. The Potato Genome Sequencing Consortium. Genome sequence and analysis of the tuber crop potato. Nature. 2011;475:189–95.

81. Sanseverino W, Hermoso A, D’Alessandro R, Vlasova A, Andolfo G, Frusciante L, et al. PRGdb 2.0: Towards a community-based database model for the analysis of R-genes in plants. Nucleic Acids Res. 2013;41:D1167–71.

82. Chen J, Hu Q, Zhang Y, Lu C, Kuang H. P-MITE: A database for plant miniature inverted-repeat transposable elements. Nucleic Acids Res. 2014;42:D1176–81.

83. Jurka J, Kapitonov V V., Pavlicek A, Klonowski P, Kohany O, Walichiewicz J. Repbase Update, a database of eukaryotic repetitive elements. Cytogenet Genome Res. 2005;110:1–4.

84. Dunn NA, Unni DR, Diesh C, Munoz-Torres M, Harris NL, Yao E, et al. Apollo: Democratizing genome annotation. PLoS Comput Biol. 2019;15:E1006790.

85. Kim S, Cheong K, Park J, Kim M-S, Kim J-H, Seo M-K, et al. TGFam-Finder: An optimal solution for target-gene family annotation in eukaryotic genomes. bioRxiv. 2018;

86. Letunic I, Doerks T, Bork P. SMART: Recent updates, new developments and status in 2015. Nucleic Acids Res. 2015;43:D257–60.

87. Lupas A, Van Dyke M, Stock J. Predicting coiled coils from protein sequences. Science (80-). 1991;252:1162–4.

88. Li L, Stoeckert CJ, Roos DS. OrthoMCL: Identification of ortholog groups for eukaryotic genomes. Genome Res. 2003;13:2178–89.

89. Katoh K, Standley DM. MAFFT multiple sequence alignment software version 7: improvements in performance and usability. Mol Biol Evol. 2013;30:772–80.

90. Kumar S, Stecher G, Tamura K. MEGA7: Molecular Evolutionary Genetics Analysis Version 7.0 for Bigger Datasets. Mol Biol Evol. 2016;33:1870–4.

91. Capella-Gutiérrez S, Silla-Martínez JM, Gabaldón T. trimAl: A tool for automated alignment trimming in large-scale phylogenetic analyses. Bioinformatics. 2009;25:1972–3.

92. Stamatakis A. RAxML version 8: A tool for phylogenetic analysis and post-analysis of large phylogenies. Bioinformatics. 2014;30:1312–3.

93. van Dongen SM. Graph clustering by flow simulation. Ph.D. Thesis. Universiteit Utrecht, Utrecht, The Netherlands; 2000.

94. Buchfink B, Xie C, Huson DH. Fast and sensitive protein alignment using DIAMOND. Nat Methods. 2014;12:59–60.

95. Löytynoja A. Phylogeny-aware alignment with PRANK. Methods Mol Biol. 2014. p. 155–70.

96. Vasimuddin M, Misra S, Li H, Aluru S. Efficient architecture-aware acceleration of BWA-MEM for multicore systems. IEEE Parallel Distrib Process Symp. 2019;

97. Li H, Handsaker B, Wysoker A, Fennell T, Ruan J, Homer N, et al. The Sequence Alignment/Map format and SAMtools. Bioinformatics. 2009;25:2078–9.

98. Murrell B, Moola S, Mabona A, Weighill T, Sheward D, Kosakovsky Pond SL, et al. FUBAR: A fast, unconstrained bayesian AppRoximation for inferring selection. Mol Biol Evol. 2013;30:1196–205.

99. Murrell B, Wertheim JO, Moola S, Weighill T, Scheffler K, Kosakovsky Pond SL. Detecting individual sites subject to episodic diversifying selection. PLoS Genet. 2012;8:e1002764.

100. Rodriguez F, Wu F, Ané C, Tanksley S, Spooner DM. Do potatoes and tomatoes have a single evolutionary history, and what proportion of the genome supports this history? BMC Evol Biol. 2009;9:191.

